# The bakers’s yeast Msh4-Msh5 associates with double-strand break hotspots and chromosome axis during meiosis to promote crossovers

**DOI:** 10.1101/2020.07.24.219295

**Authors:** Krishnaprasad G. Nandanan, Ajith V. Pankajam, Sagar Salim, Miki Shinohara, Gen Lin, Parijat Chakraborty, Lars M. Steinmetz, Akira Shinohara, Koodali T. Nishant

## Abstract

Segregation of homologous chromosomes during the first meiotic division requires at least one obligate crossover/exchange event between the homolog pairs. In the baker’s yeast *Saccharomyces cerevisiae* and mammals, the mismatch repair-related factors, Msh4-Msh5 and Mlh1-Mlh3 generate the majority of the meiotic crossovers from programmed double-strand breaks (DSBs). To understand the mechanistic role of Msh4-Msh5 in meiotic crossing over, we performed genome-wide ChIP-sequencing and cytological analysis of the Msh5 protein in cells synchronized for meiosis. We observe that Msh5 associates with DSB hotspots, chromosome axis, and centromeres. We found that the initial recruitment of Msh4-Msh5 occurs following DSB resection. A two-step Msh5 binding pattern was observed: an early weak binding at DSB hotspots followed by enhanced late binding upon the formation of double Holliday junction structures. Msh5 association with the chromosome axis is Red1 dependent, while Msh5 association with the DSB hotspots and axis is dependent on DSB formation by Spo11. Msh5 binding was enhanced at strong DSB hotspots consistent with a role for DSB frequency in promoting Msh5 binding. These data on the *in vivo* localization of Msh5 during meiosis have implications for how Msh4-Msh5 may work with other crossover and synapsis promoting factors to ensure Holliday junction resolution at the chromosome axis.

**AUTHOR SUMMARY:** During meiosis, crossovers facilitate physical linkages between homologous chromosomes that ensure their accurate segregation. Meiotic crossovers are initiated from programmed DNA double-strand breaks (DSBs). In the baker’s yeast and mammals, DSBs are repaired into crossovers primarily through a pathway involving the highly conserved mismatch repair related Msh4-Msh5 complex along with other crossover promoting factors. *In vitro* and physical studies suggest that the Msh4-Msh5 heterodimer facilitates meiotic crossover formation by stabilizing Holliday junctions. We investigated the genome-wide *in vivo* binding sites of Msh5 during meiotic progression. Msh5 was enriched at DSB hotspots, chromosome axis, and centromere sites. Our results suggest Msh5 associates with both DSB sites on the chromosomal loops and with the chromosome axis to promote crossover formation. These results on the *in vivo* dynamic localization of the Msh5 protein provide novel insights into how the Msh4-Msh5 complex may work with other crossover and synapsis promoting factors to facilitate crossover formation.

## INTRODUCTION

During meiosis in baker’s yeast, crossover formation is initiated from programmed DNA double-strand breaks (DSBs) made by Spo11 and accessory proteins [1–3]. Crossovers, together with sister chromatid cohesion, facilitate homologous chromosome segregation by physically linking the homologs and properly orienting them towards opposite poles of the bipolar spindle [4]. A fraction of the DSBs are repaired using the homologous chromosome as a template to generate crossovers, and the rest are repaired either as noncrossovers using the homolog or using the sister chromatid as template [5, 6]. In humans, failure to obtain at least one crossover per homolog pair can lead to aneuploid gametes that result in birth defects in offspring [7].

Meiotically induced DSBs are non-randomly positioned in the genome. Multiple factors, including open chromatin structure, the presence of certain histone modifications, and the binding of sequence-specific transcription factors are known to influence the spatial positioning of DSBs [8, 9]. In *S. cerevisiae*, during meiotic prophase, chromosomes are organized into a series of loops that are tethered at the base by proteinaceous axial elements mainly consisting of proteins Red1 and Hop1 and the meiotic cohesion complex containing Rec8 [10–15]. The chromatin loops originating from the cohesin enriched axis regions have conserved density (~20 loops per micron of axis length) across organisms [12, 16]. In *S. cerevisiae* loops are between 0.2 to 0.6 um long (from axis to the distal-most sequence in the loop) containing up to 20 kb of DNA [12, 17]. Within the Red1-rich axis domains, DSBs are made at a distance from the axis on the loops [16]. But DSB-promoting proteins are associated with axis regions as well, indicating that DSB formation takes place in the context of the loop-axis structure of the chromosomes [18, 19]. Based on these findings, it has been hypothesized that chromosomal loops are recruited to the proximity of the axis for DSB formation and repair [20–22]. This allows events at both loop and axis levels to be functionally linked and coordinated [16, 17]. The chromosome axis proteins influence the formation of DSBs, promote interhomolog bias, and facilitate the biased resolution of double Holliday Junction (dHJ) intermediates [23].

In *S. cerevisiae*, the major pathway for the repair of DSBs as crossovers is mediated by the ZMM group of proteins (Zip1, Zip2, Zip3, Zip4, Mer3, Msh4, Msh5, and Spo16) along with MutLγ (Mlh1 and Mlh3), Exo1, and the STR complex (Sgs1, Rmi1, Top3) to facilitate the formation of ~80% of crossovers [24–28]. The ZMM-mediated crossovers show interference and are called class I crossovers. A minor set of crossovers that are interference-independent (class II) are produced by a pathway involving the Mms4-Mus81 protein complex, which can directly act on D-loops or Holliday junction intermediates to form crossovers [29, 30].

The Msh4 and Msh5 proteins are homologs of the bacterial MutS family of mismatch repair proteins with no known function in mismatch repair [31]. The Msh4 and Msh5 proteins function together as a heterodimer [32]. In *S. cerevisiae*, both *msh4Δ* and *msh5Δ* mutants show a substantial reduction in meiotic viability along with crossover defects and Meiosis I non-disjunction [30, 33]. *In vitro* biochemical studies using purified human Msh4 and Msh5 show that these proteins bind to Holliday junction substrates and stabilize them [34]. Similarly, *in vitro* biochemical and physical studies in *S. cerevisiae* show that Msh4-Msh5 stabilizes 3’-overhangs, single-end invasion (SEI) intermediates, and Holiday junctions [24, 35]. Further, Msh4 has a degron, which is phosphorylated by cdc7 kinase to stabilize the Msh4-Msh5 complex and promote a crossover outcome [36]. Recent genetic, biochemical, and physical studies suggest that the Msh4-Msh5 complex acts in the same pathway as the Mlh1-Mlh3 complex, Exo1, and PCNA to process inter homolog joint molecule structures into crossover products [27, 37–41]. The Msh4-Msh5 complex has a role in directing the activity of the Mlh1-Mlh3 endonuclease to generate crossovers from double Holliday junctions [27, 40, 41]. In mammals, mutation of *Msh4* or *Msh5* results in sterility [42, 43]. Cell biological observations in the mouse show a large number of Msh4/Msh5 foci (~ 140 per nucleus) during an early stage (Zygotene) of meiosis [44]. As meiosis progresses, the number of Msh4/Msh5 foci decreases, and at mid pachytene stage, they are present at almost twice the number of actual crossing over sites. At this stage, almost half of the Msh4/Msh5 foci co-localize with Mlh1/Mlh3 foci, which are considered to be the sites for crossover [42, 45, 46]. Although few aspects of Msh4-Msh5 function are known, an understanding of the *in vivo* DNA binding sites, recombination substrates and localization with reference to the chromosomal organization during meiosis is lacking.

We analyzed the chromosome-binding properties of the Msh4-Msh5 complex using chromatin immunoprecipitation and sequencing (ChIP-Seq), ChIP-qPCR, and cytology in synchronized meiotic time courses. We found that Msh5 is specifically enriched at DSB hotspots [18]. Additionally, Msh5 binding is also observed at the chromosome axis and centromeres. Msh5 binding at DSB hotspots and chromosome axis is disrupted in *spo11* mutants, while axis association is disrupted in *red1*Δ. Msh5-binding sites on chromosomes are similar to Zip3 and the ZZS complex (Zip2, Zip4, Spo16) [47–49]. Further Msh5 binding at chromosome axis and centromeres also suggest crossover-independent functions related to chromosome synapsis. Our observations suggest that Msh5 is enriched at strong DSB hotspots that have high DSB frequency. These results shed novel insights into the role of the chromosome loop-axis structure on the assembly of the Msh4-Msh5 complex during meiosis. They are also useful for developing models for how Msh4-Msh5 may coordinate both crossover and synapsis events.

## RESULTS

### Genome-wide analysis shows Msh5 binding at DSB hotspots, chromosome axis, and centromeres

Msh5 protein expression starts at 2h after induction of meiosis and remains till 8h with peak expression at 5h (**Fig 1A**). To characterize the dynamic binding of Msh5 during the different stages of meiosis, we performed ChIP-Seq from meiotic extracts using an anti-Msh5 antibody. The Msh5 antibody was generated in rabbit and is polyclonal. 5ul Msh5 serum was used for each ChIP-Seq experiment. No Msh5 protein band was observed in the *msh5Δ* lysate or eluate fractions, showing the specificity of the antibody (**Fig 1B**). Msh5 ChIP-Seq was performed in quadruplicate at six-time points, from 2h to 7h at one-hour intervals. The Msh5 binding data from the four replicates (a-d) at the 5h time point was highly reproducible (r = 0.72-0.93) (**S1 A-F Fig**). Msh5 ChIP-Seq data from the four replicates were normalized using input, and genome-wide plots were generated to detect Msh5-peak positions along the chromosomes (Materials and Methods). A total of 3397 Msh5 peaks were observed from all four replicates (with p < 10^−5^, see Materials and Methods) at 5h post entry into meiosis (**S1 Table**). Previous studies of the ZMM proteins have shown that Zip3, as well as the ZZS complex (Zip2, Zip4, Spo16) mostly associate with DSB hotspots and show weak and transient association with the chromosome axis [47]. Msh5 (and Msh4) functions as a distinct complex from Zip3 and ZZS [47, 50]. We therefore analyzed Msh5 binding with reference to DSB hotspots [18] and chromosome axis sites [15]. Representative Msh5 binding plot from all four replicates for chromosome III at 3h, 4h, and 5h, shows Msh5 associates with DSB hotspots, chromosome axis sites, and also centromere position while being depleted from DSB coldspots (**S2A Fig**). The Msh5 binding sites were validated using qPCR. Binding of Msh5 at three representative DSB hotspots (*BUD23*, *ECM3*, *CCT6*), three axis sites (*Axis I*, *Axis II* and *Axis III*), two centromere locations (*CENI, CENIII*) and one coldspot (*YCRO93W*) were analyzed from the same time course in one of the replicates (d). The qPCR results confirm that Msh5 is specifically enriched at DSB hotspots, chromosome axis and centromeres relative to DSB cold spots (**Fig 1C** and **S2B Fig**). The qPCR data was further used to scale the corresponding Msh5 ChIP-Seq replicate (d) (Materials and Methods) (**S3A Fig**). From the scaled Msh5 ChIP-Seq data, maximum Msh5 binding is observed between 3-5h relative to other time points (6h, 7h) across all chromosomal locations consistent with Msh5 expression (**Fig 2A**). The scaled Msh5 ChIP-Seq plot is shown for chromosome III as a representative example (**Fig 2B**). We observed that Msh5 binds weakly to DSB hotspots at an early stage (3h) after induction of meiosis (**Fig 2B**). At later stages of meiotic progression (4h and 5h) maximum Msh5 binding was seen at DSB hotspots (**Fig 2B**). Statistical analysis of the scaled Msh5 coverage data at DSB hotspots (non-overlapping with Red1 sites) showed binding differences at 3h, 4h, and 5h are significantly different (t-test) (**Fig 2C**). Further, maximum Msh5 binding at DSB hotspots is seen at 4h −5h (**Fig 2C**). Msh5 peaks from all four replicates associated with DSB hotspots at 5h showed a median width of 0.5kb (mean 0.9kb) (**Fig 2D**). This result suggests that Msh5 binding at DSB hotspots is confined to the width of the heteroduplex region during DSB repair (1-2 kb) [51].

**Fig 1.**
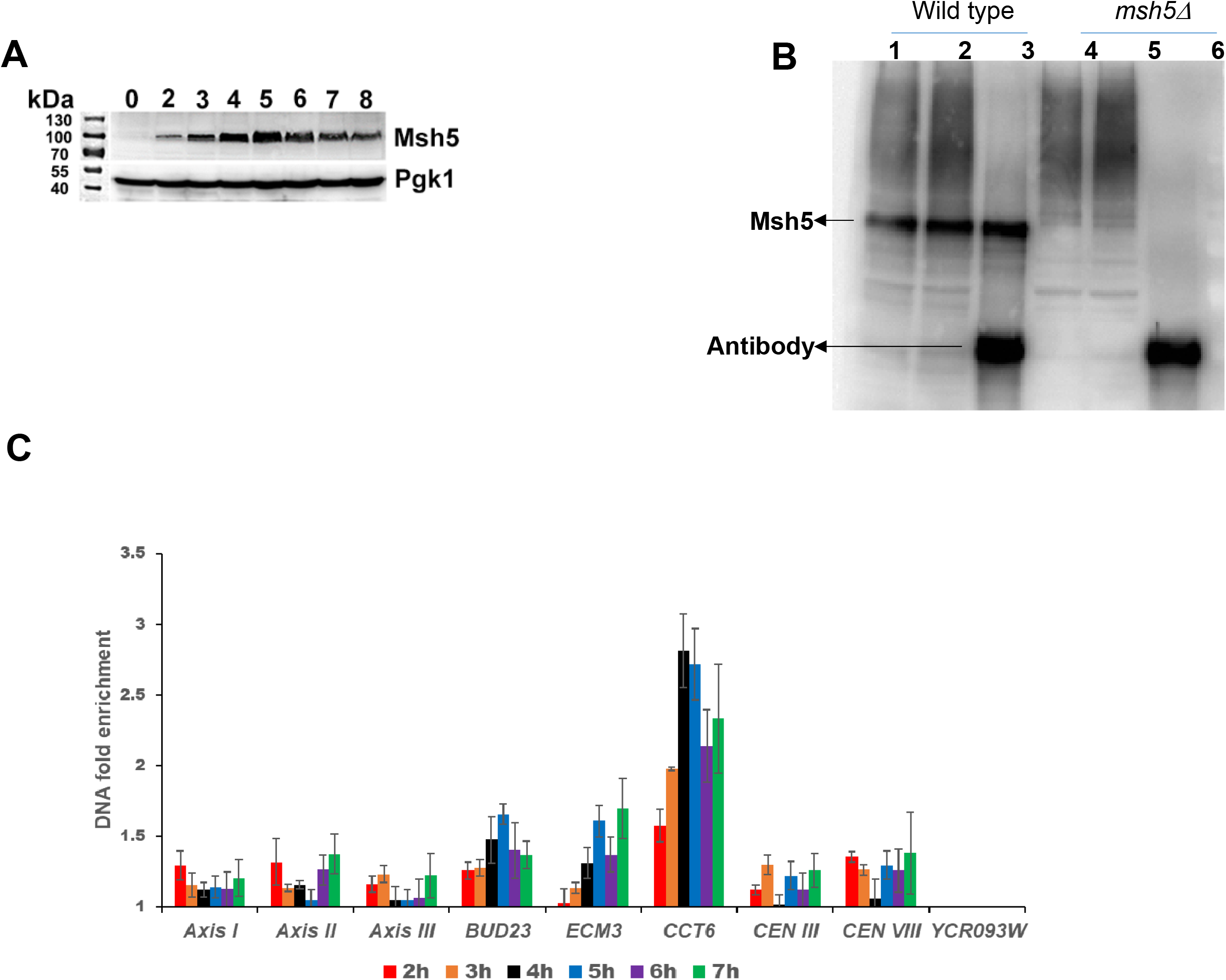
Msh5 expression and DNA binding in wild-type meiosis. **(A)** Western blot analysis of Msh5 expression pattern in synchronized wild-type meiosis. **(B)** ChIP using Msh5 antibody in wild type and *msh5Δ* strains. Lanes 1 to 3 indicate lysate, lysate after overnight incubation with Msh5 antibody and beads, and the eluate fractions in wild type. Lanes 4 to 6 indicate lysate, lysate after overnight incubation with Msh5 antibody and beads and the eluate fractions in *msh5Δ*. **C)** Msh5 enrichment at DSB hotspots (*BUD23*, *ECM3*, and *CCT6*), centromere regions (*CEN I, CEN III*) and axis regions (*AXIS I*, *AXIS II*, *AXIS III*) relative to DSB coldspot (*YCR093W*), measured using ChIP-qPCR. The samples are normalized using input. The error bars represent the standard deviation.

**Fig 2.**
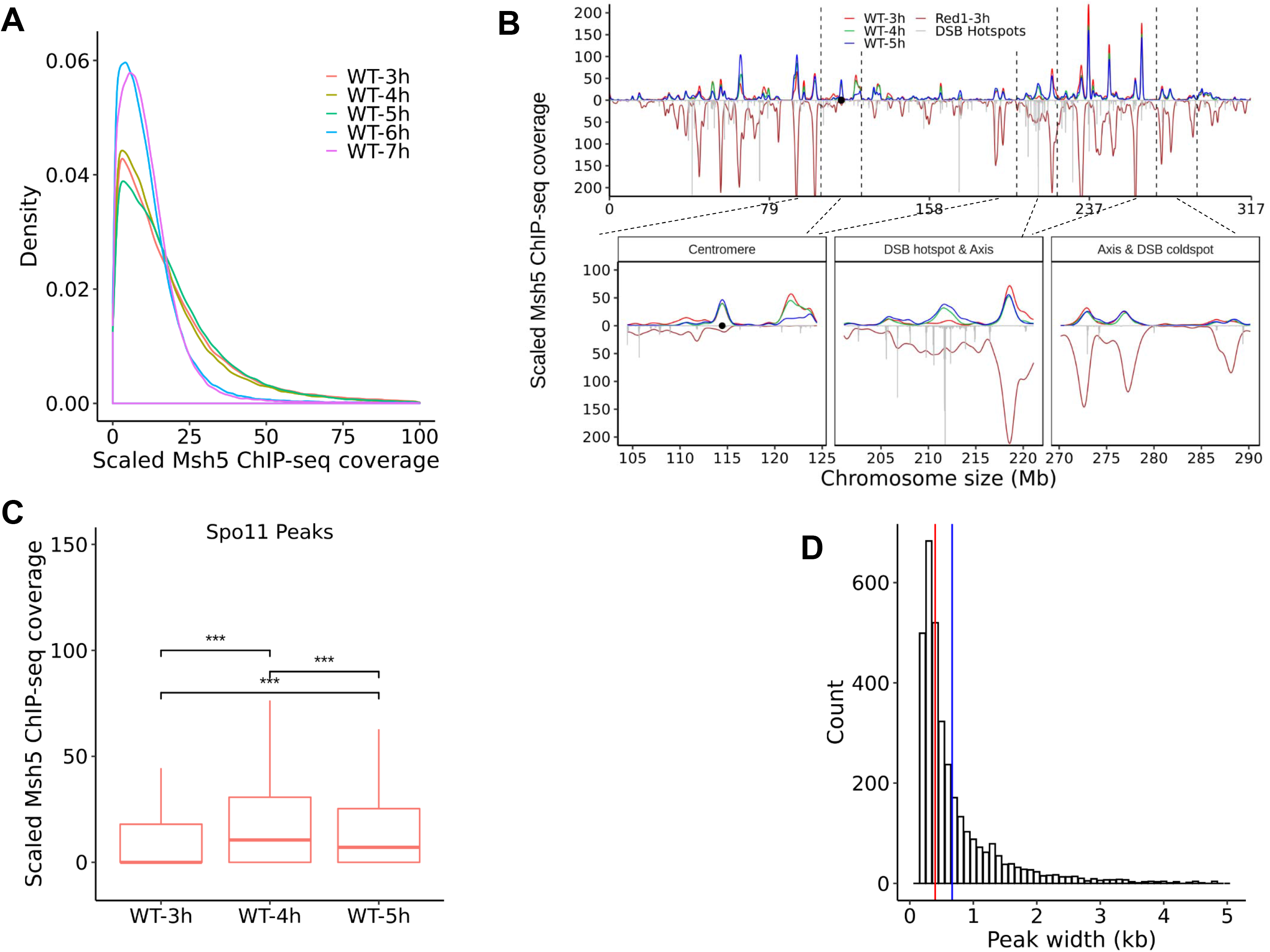
Msh5 binds at DSB hotspots, chromosome axis, and centromere positions. **A)** Density of scaled Msh5 ChIP-Seq reads at Msh5 peak locations (5h) from 3h to 7h. **B)** Representative image to show chromosomal localization of Msh5 in wild type on chromosome III (t=3, 4, and 5h after induction of meiosis) using qPCR scaled ChIP-Seq data. Zoomed in image of the centromeric region, axis region, one DSB hotspot (*BUD23*), and one DSB coldspot (*YCR093W*) is also shown. Red1 and Spo11 data are taken from, Sun et al 2015[15] and Pan et al., 2011 [18]. The black circle indicates centromere. (**C)** Coverage analysis of qPCR scaled Msh5 ChIP-Seq reads at Spo11 DSB hotspots (*** indicates p < 0.001, outliers are removed). (**D)** Width of the Msh5 binding peaks (5h time point) at DSB hotspots (not overlapping with Red1 sites) from all four Msh5 ChIP-Seq replicates.

We analyzed binding data of Msh5 in meiotic time courses at axis sites (Red1 binding regions) [15]. Representative plot for chromosome III shows the overlap of Msh5, and Red1 reads from 3h to 5h (**Fig 2B**). Msh5 binding at Red1 sites (non-overlapping with Spo11 sites) from 3h to 5h is significantly different between the time points with maximum binding at 5h (**Fig 3A**). These observations suggest Msh5 binds to both DSB hotspots and axis from 3h to 5h. Together, these observations support Msh5 association with both DSB sites at chromosomal loops and axis regions.

**Fig 3.**
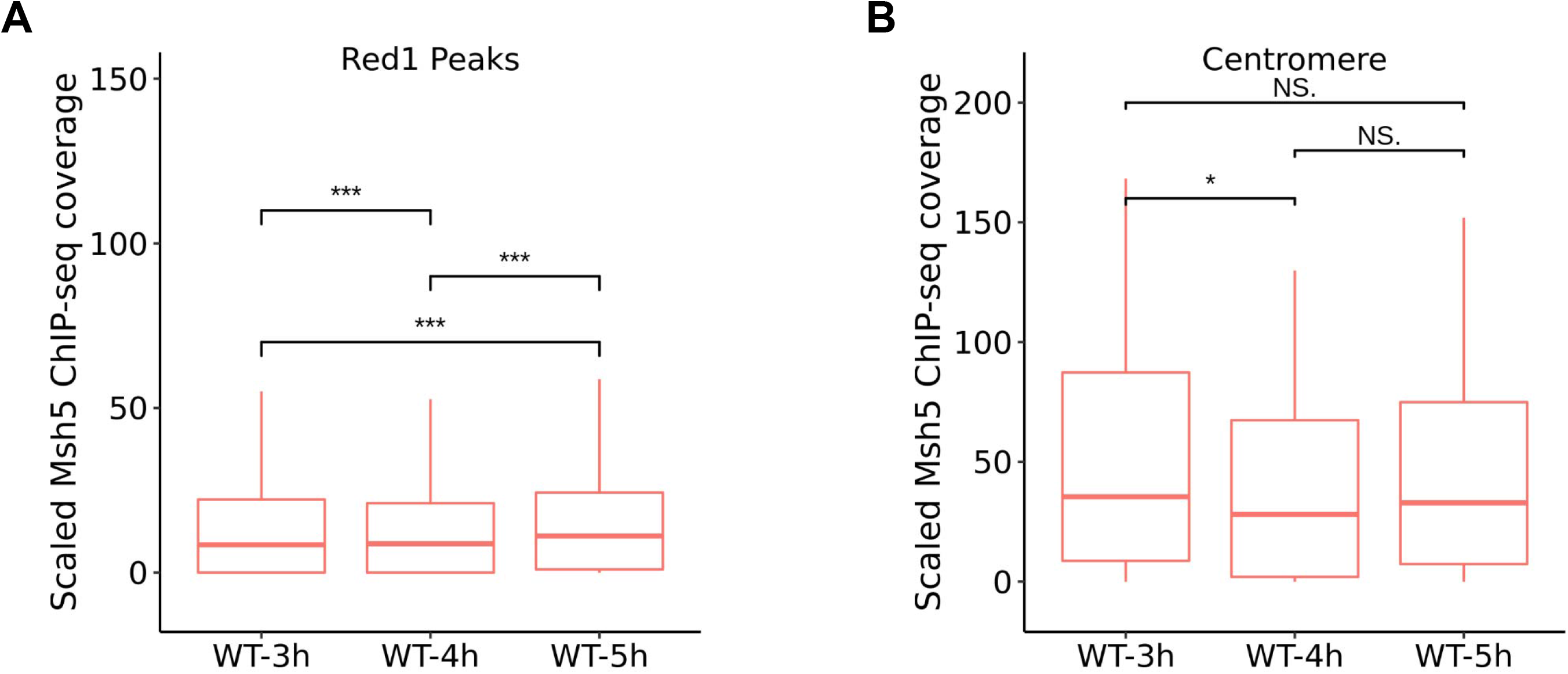
Msh5 association with chromosome axis and centromeres. **A)** Coverage analysis of qPCR scaled Msh5 ChIP-Seq reads at axis sites (Red1 peaks) (*** indicates p < 0.001, outliers are removed). **B)** Coverage analysis of qPCR scaled Msh5 ChIP-Seq reads at centromere sites (* indicates p < 0.01, outliers are removed).

In addition to DSB hotspots and axis, ChIP-qPCR and ChIP-Seq analysis also showed binding of Msh5 at representative centromere regions **(Fig 1C, 2B, S2A, B)**. Also, earlier binding of Msh5 at centromeres (e.g. at 3h) is observed compared to the DSB hotspots and axis sites (**Fig 3B**). Individual chromosome plots show Msh5 associates with centromeres on all chromosomes **(S4 Fig)**. Association with centromeres is also observed for other ZMM proteins like Zip3 [48].

qPCR analysis showed no Msh5 enrichment in *msh5Δ* mutant at DSB hotspots relative to the coldspot (**S5A Fig**). Axis and centromeres (which show reduced Msh5 enrichment compared to DSB hotspots) showed background levels of Msh5 binding in *msh5Δ*. qPCR analysis also showed no enrichment of Msh5 in *msh4Δ* mutant at DSB hotspots (**S5B Fig**). However, weak Msh5 binding is observed at axis and centromere in *msh4Δ* (**S5B Fig**) consistent with earlier cytological observations showing residual Msh5 signals in *msh4Δ* [25]. Therefore, Msh4-Msh5 complex formation is required for normal Msh5 enrichment at DSB hotspots, axis, and centromeres.

Since other ZMM proteins (Zip2, Zip3, Zip4, Spo16) have been observed to associate with the centromere, chromosome axis and DSB hotspots in ChIP-array and ChIP-Seq studies [47, 48], we re-analyzed Zip3 binding to facilitate comparison with the Msh5 binding data in the same strain background and experimental conditions. We used a Zip3 construct C-terminally tagged with three copies of the FLAG epitope [48]. The diploid strain expressing *ZIP3-His6-FLAG3* allele showed a spore viability of 97% (40 tetrads dissected), as reported previously [48]. Zip3 expression was detected using an anti-FLAG antibody (mouse) from 2h after induction of meiosis till 8h with peak expression at 5h (**S6A Fig**). Zip3 ChIP from meiotic extracts was performed at 3h, 4h, and 5h using an anti-FLAG antibody. No Zip3 protein band was observed in the untagged wild-type lysate or eluate fractions showing the specificity of the antibody (**S6B Fig**). Zip3 ChIP-Seq was performed once for 3h time point and in duplicates for 4h and 5h time points. Representative binding of Zip3 on chromosome III is shown in **S6C Fig**. Like Msh5, Zip3 shows binding to DSB hotspots, chromosome axis sites, centromere and is absent from DSB cold spots (**S6C Fig**) as observed previously [47, 48].

### Msh5 binding at DSB hotspots is stimulated by DSB repair intermediates

We performed cytology in meiotic time courses with the Msh5 antibody in several mutant backgrounds that affect DSB formation and processing to determine genetic requirements for Msh4-Msh5 localization at DSB hotspots. In wild-type cells, we observed a peak in the average number of Msh5 foci/cell (48 ± 6.5) at 5h (**Fig 4A, C**). Previous studies have also observed a similar number of Msh5 foci in wild type [36, 49]. In a *spo11*Δ mutant where DSBs are not formed, cytological observations showed no Msh5 foci (**Fig 4B, C**) at any of the time points in meiosis. Consistent with previous studies, these observations suggest that Msh5 binding to the DNA is dependent on DSB formation [49]. We asked if DSB formation and resection are sufficient for the binding of Msh5 protein. In *dmc1*Δ mutant, cells arrest after DSB formation and resection [52]. The joint molecule formation is blocked completely in *dmc1Δ* mutant [53]. These cells cannot repair DSBs because of the loss of strand exchange activity [52, 54]. Compared to wild type (48 ± 6.5 Msh5 foci), on average 22 ± 3.8 Msh5 foci were observed at 5h in a *dmc1*Δ mutant **(Fig 4A-C)**. These results indicate that an event prior to the Dmc1 function, such as DSB resection, promotes Msh5 binding. Such an observation is also consistent with *in vitro* data showing Msh4-Msh5 can bind to 3’ overhangs [35]. It has been shown that in the *dmc1*Δ mutant, the DSBs appear at the normal time but persist longer, and the cells get arrested at this stage [52]. The reduced number of Msh5 foci (22 ± 3.8) in *dmc1*Δ mutant compared to wild type (48 ± 6.5, p < 0.001) is therefore not due to reduced DSBs. Instead, this observation suggests that even though resected DSBs can recruit Msh5, the binding efficiency to resected DSB structures is reduced. Dmc1-dependent strand invasion steps, therefore, promote stable binding of Msh5.

**Fig 4.**
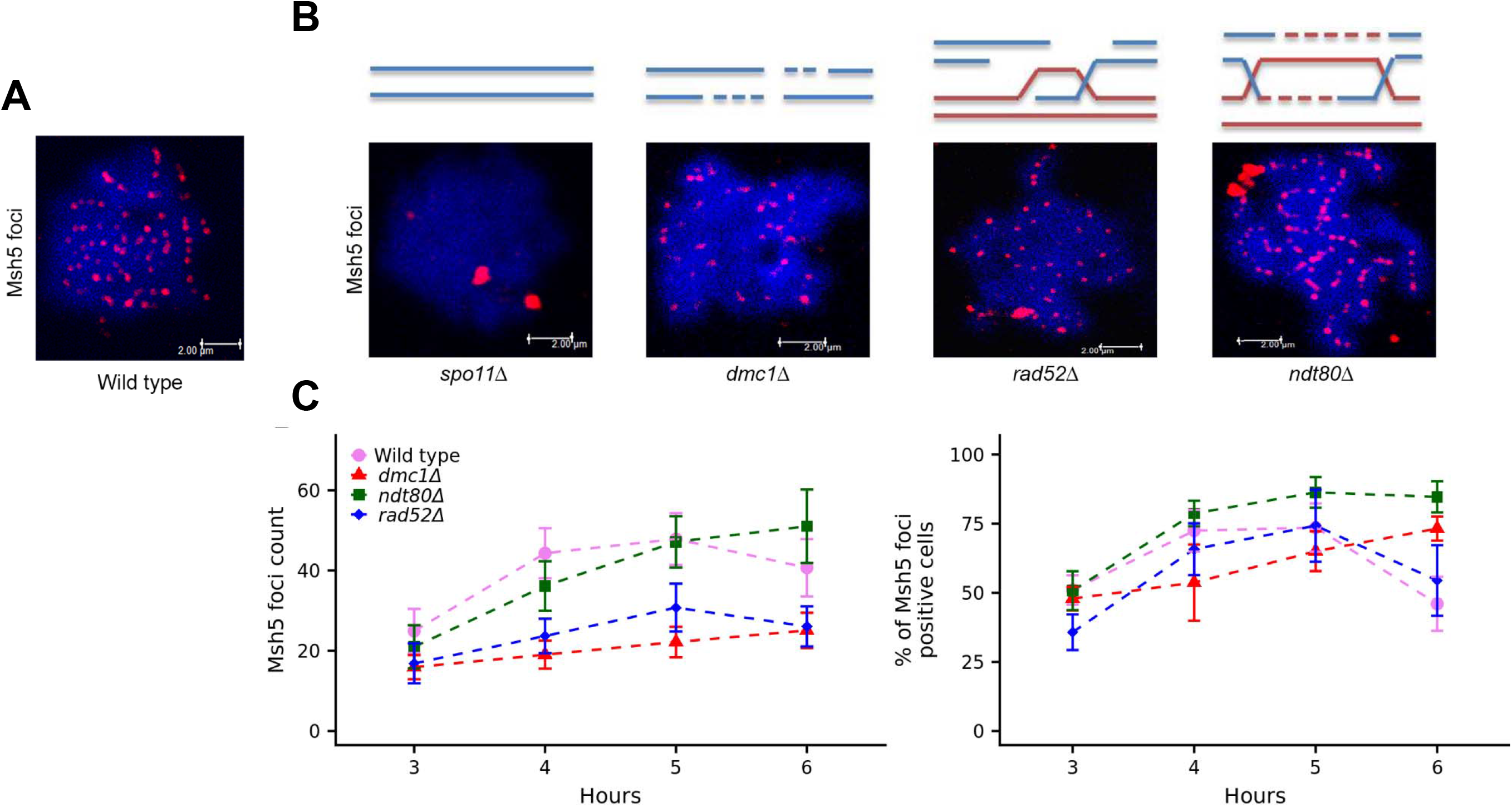
Msh5 binding is stimulated by meiotic DSB repair intermediates. **A)** Representative image of a spread meiotic nucleus from wild-type strain (t=5h) immunostained for Msh5 (red foci). **B)** Representative image of a spread meiotic nucleus from *spo11Δ, dmc1Δ, rad52*Δ, and *ndt80*Δ mutants (t=5h) immunostained for Msh5 (red foci). *spo11Δ* indicates the specificity of the Msh5 antibody. (**C)** Kinetics of Msh5 foci and the percentage of meiotic nuclei positive for Msh5 foci in synchronized meiosis in wild type and *dmc1Δ*, *rad52Δ, ndt80*Δ mutants. Approximately 50 nuclei, each from two independent experiments, were assessed for each time point (except *spo11Δ*, see Materials and Methods). The error bars represent the standard deviation.

We tested Msh5 binding in a *rad52*Δ mutant where cells efficiently proceed to strand invasion and make wild-type level of SEI intermediates, but dHJ formation is significantly reduced [55, 56]. The timing and frequency of DSBs are similar in *rad52*Δ and wild type [55]. We observed an average of 31 ± 6 Msh5 foci at 5h in the *rad52*Δ mutant (**Fig 4B, C**), which is less compared to wild type (p < 0.001). Since SEIs are similar in *rad52*Δ and wild type, but SEI + dHJs are about two times more in wild type, the number of Msh5 foci in *rad52*Δ (31 ± 6) and wild type (48 ± 6.5) suggests that SEI intermediates are efficient for recruiting Msh4-Msh5. Taken together with previous observations of lack of Msh5 foci in *spo11*Δ and *rad50S* mutants [49], our results in *dmc1Δ* and *rad52Δ* suggest in *S. cerevisiae* Msh4-Msh5 recruitment *in vivo* happens after DSB resection, which is consistent with *in vitro* biochemical data showing binding of *S. cerevisiae* and human Msh4-Msh5 at SEI structures [34, 35].

Next, we asked if dHJs stimulate maximum Msh5 binding using the *ndt80*Δ mutant that accumulates joint molecules, but the dHJs are not resolved [5]. An average of 47 ± 6.4 Msh5 foci is observed at 5h in *ndt80*Δ mutant which is similar to the maximum number of Msh5 foci seen in the wild type at 5h (p = 0.44, **Fig 4A-C**) indicating that the SEI and dHJ structures together facilitate maximum binding of the Msh4-Msh5 complex to the DNA. These results suggest that even though resected DSBs can initiate binding of the Msh4-Msh5 complex, maximum binding is achieved with joint molecule structures comprising SEIs and dHJs (see discussion). In addition, the average Msh5 foci count (51) at 6h in *ndt80Δ* is higher than wild type (40.7) due to the accumulated levels of SEIs and dHJ structures. Msh5 aggregates were observed in all the above mutants (*spo11Δ*, *dmc1Δ*, *rad52Δ*, *ndt80Δ*) and is quantified in **S7 Fig**.

### Msh5 binding depends on DSB strength

We tested the correlation between DSB frequency and Msh5 association with chromosomes. ChIP-qPCR analysis corroborated cytological observations of the absence of Msh5 binding in *spo11*Δ. ChIP-qPCR analysis showed no enrichment of Msh5 at DSB hotspots and axis relative to DSB coldspot (**S5C Fig**). But early binding (3h) at centromeres was observed, suggesting Msh5 enrichment at DSB hotspots and chromosome axis is dependent on DSB formation (**S5C Fig**). In budding yeast 3600 DSB hotspots have been identified in high resolution [18]. These DSB hotspot positions (Spo11 cutting sites) were obtained from Pan et al., 2011[18]. We defined DSB hotspots that show Msh5 peak at 5h time point (p < 10^−5^) in at least one of the four replicates as Msh5 enriched DSB hotspots. DSB hotspots that did not show Msh5 peak (or p > 10^−5^) in any of the four replicates were defined as Msh5 depleted DSB hotspots. Our analysis suggests that about half of the 3600 identified DSB hotspots (1734/3600) are enriched for Msh5 protein (**S2 Table**). We observed that Msh5 enriched DSB hotspots showed higher Spo11 oligo counts (**Fig 5A, S2 Table**). Among the top 10% of DSB hotspots (based on Spo11 oligo counts,[18]), 96% were Msh5 enriched DSB hotspots and 4% were Msh5 depleted DSB hotspots (p < 0.01) (**S2 Table**). Among the bottom 10% of DSB hotspots, 27% were Msh5 enriched DSB hotspots, and 73% were Msh5 depleted DSB hotspots (p <0.01). This observation suggests Msh5 is easily detected at active DSB hotspots as more cells in the population would have the break.

**Fig 5.**
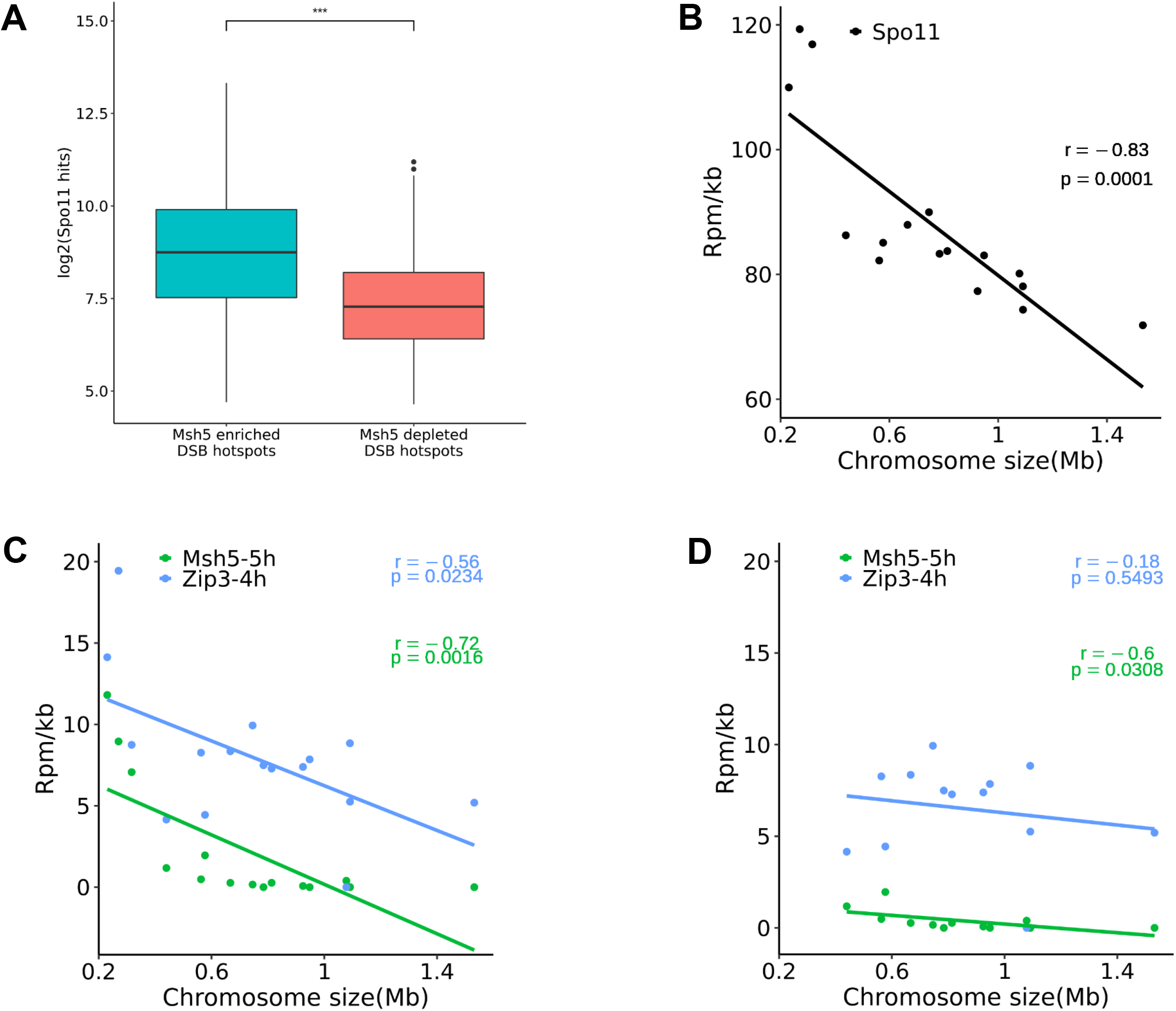
Msh5 binding at DSB hotspots is dependent on DSB frequency. **(A)** Log2 of Spo11 reads [18] at Msh5 enriched and depleted DSB hotspots. **B)** Spo11 peak density plotted as a function of chromosome size [18]. **C)** Msh5 and Zip3 read density plotted as a function of chromosome size. **D)** Msh5 and Zip3 read density as a function of chromosome size, excluding the shortest chromosomes (I, III and VI).

Small chromosomes have a higher density of crossovers than long chromosomes, which is correlated with a higher Spo11 oligo density on the smaller chromosomes [18, **Fig 5B**]. Msh5 read density (reads / Kb) analyzed from all four replicates was negatively correlated with chromosome size (r = −0.72; p = 0.0016) consistent with more crossovers / kb and higher DSB activity on smaller chromosomes (**Fig 5C**). These observations also support Msh5 enrichment is dependent on DSB activity. Zip3 read density showed weaker correlation with chromosome size (r = −0.56, p = 0.02) (**Fig 5C**). A recent study shows that the three smallest chromosomes (I, III, and VI) have unusually high DSB density due to distinct regulation of DSB formation that ensures their accurate segregation [57]. Although Spo11 oligo density (and crossover density) is negatively correlated with chromosome size even after excluding the three smallest chromosomes [57], Msh5 read density showed weak negative correlation (r = −0.6, p = 0.03), while Zip3 read density (r = −0.18, p = 0.55) was uncorrelated with chromosome size (**Fig 5D**).

### Msh5 binding to the chromosome axis is disrupted in *red1*Δ mutant

Since Msh5 binding is positively correlated with the Red1 axis associated sites, we tested whether loading of Msh5 is dependent on the Red1 protein. Red1, an axial element protein, is essential for normal DSB formation by recruiting DSB factors to the axis, synapsis, and for promoting interhomolog DSB repair through its role in meiotic checkpoint signaling [19, 58–61]. Cytological observations showed that the number of Msh5 foci in *red1*Δ was significantly less than the number of Msh5 foci in wild type at all time points (3h, 4h, 5h, 6h) in meiotic time courses with no overlap in the standard deviation (**Fig 6A, S1 Data**). At 5h an average of 16 ± 5.3 Msh5 foci is observed in *red1*Δ compared to 48 ± 6.5 foci (p < 0.001) in wild type (**Fig 6A**). Since the number of Msh5 foci is less compared to wild type, we also examined the DSB frequency using Rad51 foci counts. We observed an average of 20 ± 4.3 Rad51 foci at a three-hour time point when the numbers of DSBs are at peak level (**Fig 6B**). Previous studies have also shown that DSBs are decreased in *red1Δ* mutant [60]. The average number of Rad51 foci in *red1*Δ is significantly less compared to the number of Rad51 foci in wild type at 3h (49 ± 15, p <0.001) [58]. The reduced number of Msh5 foci in *red1*Δ is most likely due to overall reduction in DSB levels, and since DSB repair is mostly off the sister chromatid [61–63].

**Fig 6.**
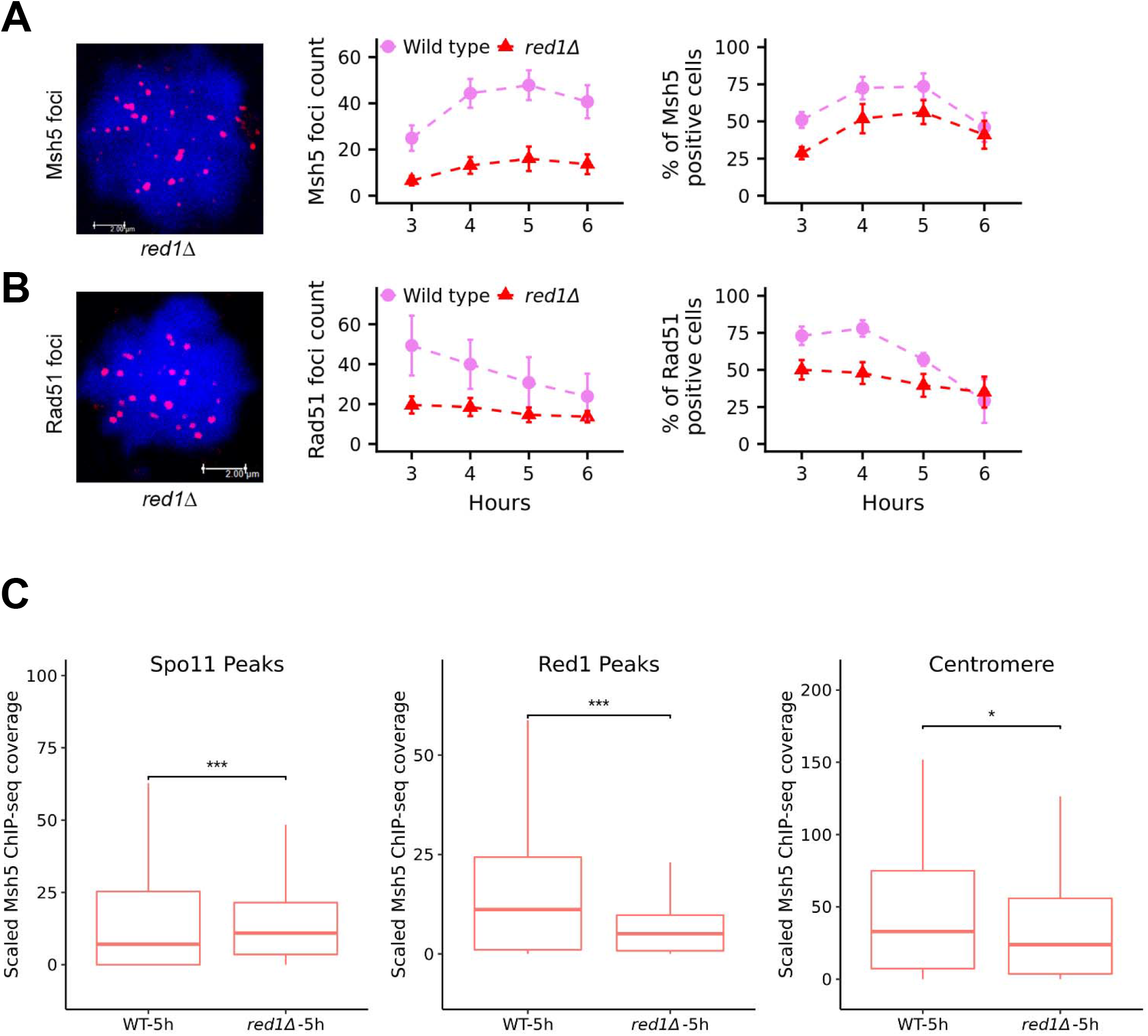
Msh5 association with chromosomes in *red1*Δ mutant. Representative image of **A)** Msh5 (t = 5h) and **B)** Rad51 (t = 3h) foci and kinetics of its accumulation in *red1Δ* mutant. Approximately 50 nuclei, each from two independent experiments, were assessed for each time point. The error bars represent the standard deviation. **(C)** Coverage analysis of qPCR scaled Msh5 ChIP-Seq reads at Spo11 DSB hotspots, Red1 peaks, and centromere positions (*** indicates p < 0.001 and * indicates p < 0.01, outliers are removed).

ChIP-qPCR analysis showed no enrichment of Msh5 at representative axis sites relative to the DSB coldspot in *red1*Δ at an early time point (3h) (**S5D Fig**). Msh5 association was maintained at representative DSB hotspots and centromeres (**S5D Fig**). ChIP-Seq analysis of Msh5 protein in the *red1*Δ mutant scaled with qPCR data (at 5h, **S3B Fig**) confirmed Msh5 is depleted at axis sites and maintained at DSB hotspots and centromere sites (**Fig 6C**). Overall these observations suggest that Red1 is essential for Msh5 binding at axis regions but not at DSB sites and centromere regions.

## DISCUSSION

Previous studies using cytological, biochemical, and physical approaches in various organisms have given insights into the timing of Msh4/Msh5 action, and it’s *in vitro* DNA binding properties. Purified human Msh4-Msh5 has a high affinity towards SEI and dHJs [34], and the *S. cerevisiae* Msh4-Msh5 has also been shown to bind 3’ overhangs, pro-Holliday junctions and Holliday junction structures [34, 35]. But the genome-wide dynamic localization of the Msh4-Msh5 complex with reference to its *in vivo* binding sites, meiotic chromosomal organization, and mutations in other meiotic recombination genes is not well understood. The current model of meiotic recombination based on *in vitro* biochemical analysis suggests Msh5 associates with 3’ overhangs, SEI, and dHJs. We demonstrate that *in vivo*, Msh5 associates with DSB hotspots, chromosome axis, and centromere sites. Recruitment of Msh5 to DSB hotspots and chromosome axis requires DSB formation and is observed as early as the stage of DSB resection (**Fig 4**).

Msh5 foci in *dmc1*Δ mutants, where DSBs are formed and resected but cannot undergo strand invasion is consistent with *in vitro* studies (**Fig 4**) [35]. It is possible that Msh5 binding in *dmc1Δ* may reflect aberrant binding due to the hyperresected DSBs. Further studies are required to explain whether this binding has any functional relevance or is just related to its recruitment. Although resected DSBs are sufficient for the initial recruitment of Msh5, the optimal binding of Msh5 requires the formation of dHJs (**Fig 4**). ChIP-Seq results also suggest weak binding of Msh5 at DSB hotspots at the early stages of meiosis (3h) and maximal binding at DSB hotspots around 4-5h that corresponds to joint molecule formation. These results, along with previous studies showing Msh2 associates with DSBs *in vivo* [64] suggest a general property of MSH proteins to bind DNA that facilitates specific binding to their major substrates.

Msh5 binding is observed at chromosome axis sites from 3 to 5h with maximum binding at 5h. This observation supports previous cytological studies that show localization of Msh5 and Msh4 at the chromosome axis [50, 65]. The observation that Msh5 associates with both DSB and chromosome axis sites suggest that Msh5 may participate in both the processing of recombination intermediates and the assembly of the synaptonemal complex like some of the other ZMM proteins [66]. Thus it may have a role in crossover formation and synapsis. It is also possible that Msh5 binding to the axis is indirect via interactions with axis proteins. A recent study has implicated interactions between Msh5 and Red1 [47].

The Msh5 protein also shows binding to centromere sites on all chromosomes. Other ZMM proteins like Zip2, Zip3, Zip4, and Spo16 also show centromere localization. It is suggested that Zip1, Zip2, Zip3, and Spo16 binding to centromeres promotes synapsis initiation from centromeres, besides crossover sites [67]. It is also suggested that Zip1 localization to centromere facilitates non-exchange chromosome segregation. Just as Msh5 association with chromosome axis, may facilitate synapsis, it is possible that Msh5 localization to centromeres may also facilitate synapsis initiation, especially as it is observed even in *spo11*Δ mutant at early time points (**S5C Fig**). Alternatively, it is possible that Msh5 binding at centromere may reflect a docking site for the protein or is indirect through interactions with other ZMM or centromeric proteins.

Cytological and ChIP-array studies also show that some of the ZMM proteins Zip1 and Zip3 associate with the chromosome axis [19, 48]. A recent study by De Muyt et al. 2018 [47] suggests that ZZS (Zip2, Zip4, Spo16) and Zip3 show strong association with DSB hotspots and weak and transient association with chromosome axis. Similar to Zip3 and the ZZS complex, which associate with the centromere, chromosome axis, and DSB hotspots in a dynamic manner [48], Msh5 associates with centromeres, chromosome axis and DSB hotspots (**Fig 1C**, **Fig 2B**). Further, Msh5 loading is dependent on Zip3 protein [25], and Zip3 and Msh4 foci co-localize cytologicaly. This is supported by our observation that Zip3 and Msh5 occupy similar chromosomal features (**S6C Fig**, **Fig 1C**, **Fig 2B**). Although there are similarities between Zip3 and Msh5 binding sites, Msh5 binding is also stimulated by DSB frequency (**Fig 5**, **S2 Table**). These observations support a direct role for Msh5 in the mechanism of crossover formation.

### Model for Msh4-Msh5 function in crossing over

Chromosome domains with high DSB activity are enriched for Red1 at the axis. But within the Red1 enriched domain, there is a negative correlation between Spo11/Dmc1 and Red1 sites since DSBs occur on the loop while Red1 sites are on the axis [16, 19]. We observed that about half of all DSB hotspots are detectably enriched for Msh5. The Msh5 localization supports the model that recombination occurs in sequences that are at a distance from the base of the chromosome loops [16]. The Msh5 binding data also suggests that within axis protein-enriched regions, the Msh4-Msh5 complex binds to the DSB hotspots that are known to occur at a distance from the axis. We also observe Msh5 association with the chromosome axis from 3h to 5h time point. The axial association is absent in a *spo11*Δ mutant (**S5C Fig**), which suggests, DSBs may promote Msh5 assembly on the chromosome axis. At 4-5h, Msh5 binds both DSB and axis optimally, which may facilitate communication between DSB sites and axis through Msh5 for crossover formation. Msh5 may be part of the recombinosome for loops tethering to the axis. Based on our observations that the Msh4-Msh5 complex binds to DSB hotspots on chromosomal loops and also on the axis, a model for how Msh4-Msh5 may work in crossover formation is shown in **Fig 7**.

**Fig 7.**
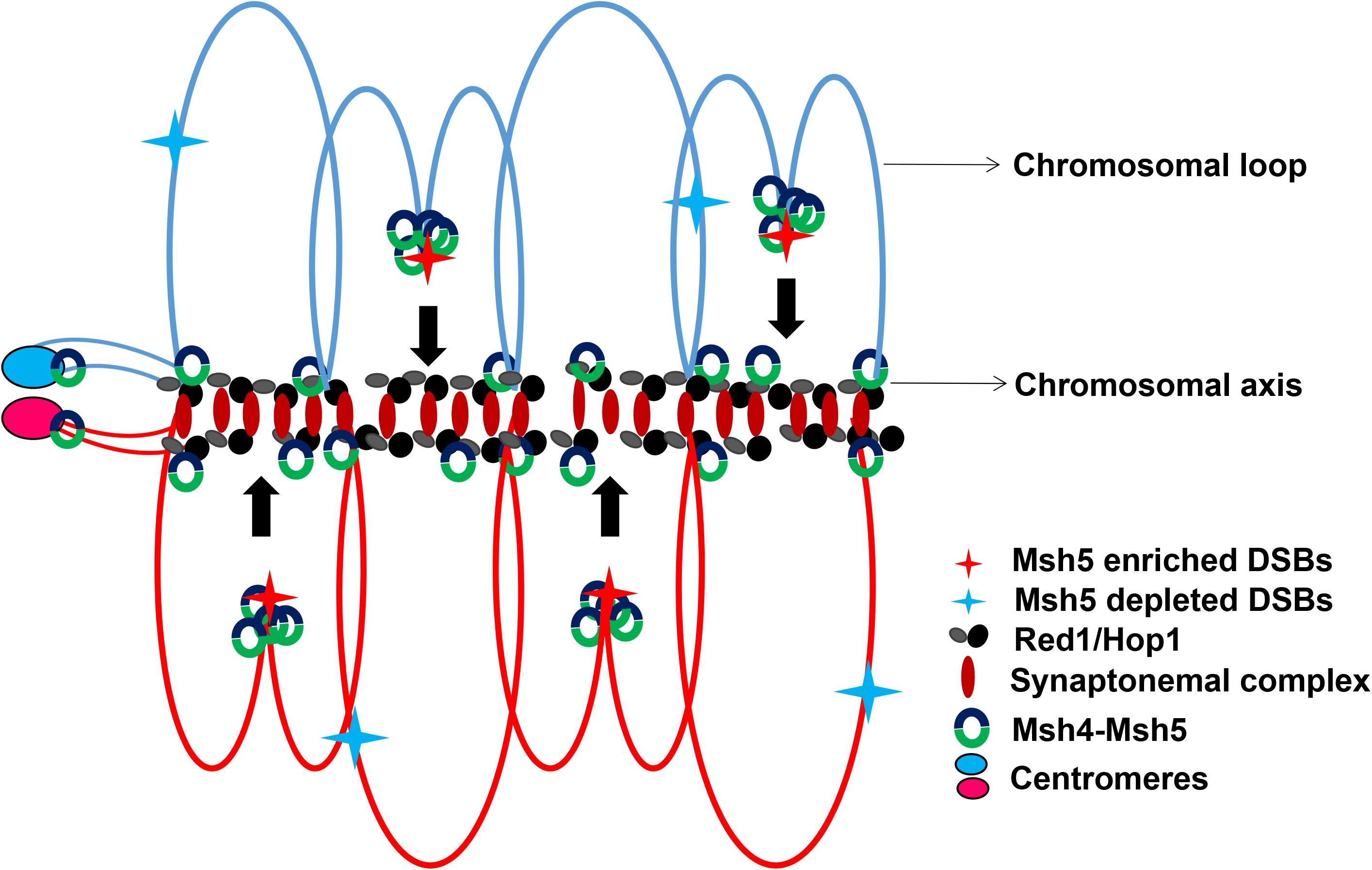
Model for the chromosomal localization of the Msh4-Msh5 complex during meiosis. During meiosis binding of the axis proteins (Red1/Hop1) define the chromosomal axis region with intervening loop regions. The loops are recruited to the axis region for DSB formation. Msh4-Msh5 associates with the DSB hotspots and axis after DSB formation at early stages (3h) of meiosis. At later stages (4-5h), enhanced binding of Msh5 is observed at DSB hotspots consistent with the generation of joint molecule structures that promote Msh5 binding. Also, Msh5 is enriched at strong DSB hotspots. Msh5 binding to both loop and axis regions may promote loop-axis interaction for crossover formation.

Similar to the loop – axis interaction required for meiotic DSB formation, we speculate a loop-axis interaction model for crossover formation. Specifically, Msh4-Msh5 association on chromosome loops and axis following DSB formation may facilitate the recruitment of the Mlh1-Mlh3 complex at these locations. Recent *in vitro* studies suggest that the Mlh1-Mlh3 complex forms unidirectional polymers on DNA and that polymerization is essential for its nicking activity [27, 68]. Since Msh4-Msh5 encircles both interacting duplexes during joint molecule formation, it could direct polymerization on both duplex arms. The Mlh1-Mlh3 complex could polymerize directionally from the loop associated Msh4-Msh5 complex towards the axis. Since the Msh4-Msh5 complex is also associated with the axis, it could facilitate loop-axis interaction for crossover formation near the axis by Mlh1-Mlh3. Also, human Msh4-Msh5 is known to stimulate human Mlh1-Mlh3 endonuclease activity [40, 41]. Crossover formation at the axis can facilitate long-range interference through the dissipation of mechanical stress along the axis [69, 70]. This axis-loop interaction during crossover formation may also be required to convert crossovers into chiasma. In conclusion, these data shed novel insights into the *in vivo* assembly and functions of the Msh4-Msh5 complex in relation to meiotic chromosome organization and DSB activity.

## MATERIALS AND METHODS

### Yeast strains and media

Yeast strains used in this study are derivatives of *S. cerevisiae* SK1 strain and are listed in the **S3 Table**. Yeast strains were grown on either yeast extract–peptone–dextrose (YPD) or synthetic complete media at 30^0^ [71–73]. Sporulation medium was prepared as described in Argueso et al. (2004) [30]. Mutants were generated by direct transformation using standard techniques [74]. When required, the drugs geneticin (Invitrogen), nourseothricin (Werner BioAgents, Germany), and hygromycin (Sigma) were added to the media at prescribed concentrations [75].

### Meiotic synchronization

Meiotic synchronization was performed using SPS method as described [76]. In short, diploid colonies were inoculated in 5 ml YPD medium and grown for 24h at 30° C to reach saturation. From the saturated YPD culture ~ 5 x 10^6^ cells/ml were inoculated to liquid SPS (0.5% yeast extract, 1% peptone, and 0.67% yeast nitrogen base without amino acids, 1% potassium acetate, 0.05M potassium biphthalate, and pH 5.5) medium and grown for 7h at 30° C. This culture was used to inoculate 500 ml of liquid SPS at cell density ~ 3 x 10^5^ cells/ml and grown for 12-16h at 30° C until the cell density reached ~4 x 10^7^ cells/ml. Cells were collected by centrifugation, washed twice with 1% potassium acetate and re-suspended in 400 ml of sporulation (2% Potassium Acetate) medium at cell density ~ 4 x 10^7^ cells / ml and incubated at 30° C with shaking. Meiotic progression was monitored by examining the nuclear divisions using DAPI staining.

### Chromatin Immunoprecipitation

Msh5 ChIP was performed as described in [19, 76, 77] with minor modifications, using native Msh5 antibody and Protein A Sepharose beads (GE Healthcare). Briefly, from a synchronous meiotic culture, 50 ml of samples were collected at various time points (as described in the results) and crosslinked using formaldehyde. Cells were lysed using Beadbeater. The cell extract was sonicated using Bioruptor (Diagenode) and the debris were removed by centrifugation. 50 ul of Protein A sepharose beads were used to preclear the cell lysate from each sample. To the pre-cleared lysate, Msh5 antibody and Protein A sepharose beads blocked with BSA was added. After incubation, lysate was removed by centrifugation and beads were washed to remove non-specific binding. The immunoprecipitated material was eluted from the beads and decrosslinked along with RNAse A treatment. The DNA was purified following Proteinase K treatment using QIAGEN PCR purification kit. Lysate samples were collected before and after the immunoprecipitation and after elution to check for IP efficiency by Western blotting. Zip3 ChIP was performed using anti-FLAG antibody and Protein G Dynabeads (Life technologies). For Msh5 western blot, anti-Msh5 antibody was used at 1:4000 dilution, and anti-rabbit HRP secondary antibody was used at 1:10000 dilution. For Zip3 western blot, anti-FLAG antibody was used at 1:4000 dilution, and anti-mouse HRP secondary antibody was used at 1:10000 dilution. Anti-PGK1 antibody was used at 1:30000 dilution, and anti-mouse HRP secondary antibody was used at 1:40000 dilution. The immunoprecipitated DNA was used for sequencing as well as qPCR. For wild type, four ChIP samples were sequenced for each time point (2-7h). For the *red1*Δ mutant, three ChIP samples were sequenced at the indicated time points. The DNA fragments were sequenced on Illumina HiSeq 2500/4000 platform at Fasteris, Switzerland. qPCR was performed on a fraction of the DNA sample used for ChIP-Seq for wild type and mutants. The DNA enrichment of the ChIP samples were calculated with respect to the input for wild type and mutants at each time point from three technical replicates.

### Cytology

Yeast chromosome spreads were prepared as described [25, 56] and stained with anti-Msh5 and anti-Rad51 antibody (Rabbit polyclonal antibody from Akira Shinohara) at 1: 500 dilution followed by anti-Rabbit TRITC conjugated secondary antibody (Jackson ImmunoResearch) at 1: 1500 dilution. Immunofluorescence images were captured using a 63x oil immersion objective on a Leica SP5 confocal microscope. For each time point, approximately 50 images were analyzed from two independent experiments. For *spo11Δ*, only 30-50 images were analyzed at each time point from a single experiment as previous studies show the absence of Msh5 foci in *spo11Δ*. Image analysis and 3D deconvolution were done using Leica Application Suite (LAS) Advanced Fluorescence (AF) Lite 2.8.0 software. All foci counts are shown as mean ± standard deviation.

### Bioinformatic analysis of Illumina sequence data

Quality and statistics of raw reads were analyzed using FastQC (version 0.11.5). Preprocessing was decided based on the FastQC report summary. Removal of Illumina adapters and trimming of raw sequence reads was done using Trimmomatic (version 0.36). Processed reads (~2-8 Million reads per samples) were aligned to the S288c reference genome (version R64-1-1) using bowtie2 (version 2.3.0) since the SK1 reference genome assembly is incomplete. Previous studies have used both S288c and SK1 as reference genomes and showed that the results are similar [15, 18]. Samples having more than 75% alignment rate and with more than 2 million aligned reads were used for the downstream analysis. Statistics of the mapped reads generated from the alignment program were analyzed for uniquely mapped reads as well as for multiple mapped reads. Samples having an alignment rate of less than 30% and less than 0.5 million reads were discarded from the analysis. Unmapped reads were analyzed using blast to detect any type of contamination in the sample. After analyzing the alignment statistics, conversion of aligned file format (SAM) to its compressed file format (BAM), indexing and sorting was done using samtools (version 1.3.1). All the downstream analyses were done using R (version 3.3). All raw sequence data for this study are available from the National Centre for Biotechnology Information Sequence Read Archive under accession number SRP129066.

### Genome-wide profile normalization using NCIS

The S288c reference genome was partitioned into equal-sized bins (200 bp), and reads per bin were calculated from the samples. Reads that are mapped to N locations were assigned a score of 1/N for each read in case of multi-mapped reads. Normalization was performed in order to account for factors affecting the data distribution, such as an error during library preparation, sequencing, and sample preparation. For this study, we performed normalizations with the control sample (Input for Msh5 ChIP and wild type untagged for Zip3 ChIP) using NCIS (Normalization of ChIP-seq). NCIS estimates the fraction of background from the control sample using a data-adaptive length of the window and a data-adaptive threshold assuming that background tends to have lower counts than enriched regions [78]. After normalizing the counts using NCIS, backgrounds were subtracted from their respective control samples and averaged, followed by genome-wide smoothening by ksmooth with a bandwidth of 1 kb.

### Peak calling using MACS

To identify the Msh5 peaks, only reads that are uniquely aligned were taken into account. MACS (Model-based Analysis for ChIP-Seq) (http://liulab.dfci.harvard.edu/MACS/) [79] was used to identify peaks from the sample. MACS uses a dynamic Poisson distribution to identify local biases in peak detection. Msh5 peaks were called from the pooled set of four replicates. Msh5 peaks with a p-value greater than 10^−5^ were filtered out, and the final set of Msh5 peaks is shown in **S1 Table**. Spo11 DSB hotspots (non-overlapping with Red1 sites) and axis sites (Red1 bound regions non-overlapping with Spo11 sites) were considered as Msh5 enriched if the Msh5 peak showed minimum one base pair overlap with the Spo11 hotspot or the Red1 peak respectively (**S2 Table**). The raw data for calling Red1 peaks were extracted from Sun et al., 2015[15]. Red1 peaks were called following the same criteria described for calling Msh5 peaks, except for p-value, which was set at less than 10^−15^ as per Sun et al., 2015[15]. Zip3 reads were analyzed similarly to Msh5 reads.

### ChIP-qPCR scaling of Msh5 ChIP-Seq reads

For wild type, one of the four Msh5 Chip-Seq replicates was scaled with the corresponding qPCR data. We performed normalization of the Msh5 Chip-Seq data with respect to its input using NCIS [78]. Linear regression model was generated between the normalized Msh5 Chip-Seq data and the qPCR data [**S3A Fig**] using R scripts provided by Hajime Murakami and Scott Keeney [80]. The linear regression model was used to scale the Msh5 ChIP-Seq data. A similar process was used to scale Msh5 ChIP-Seq reads in the *red1*Δ mutant.

### Statistical tests

For all count data (foci, box plots) student’s t-test was used to test for statistical significance. For all correlation analysis, Pearson’s correlation coefficient was used. Numerical data underlying all graphs are shown in **S1 Data**.

## ACKNOWLEDGEMENTS

We thank Valerie Borde for the strain with Zip3-FLAG construct and Scott Keeney and Hajime Murakami for the R script for scaling ChIP-Seq data. We thank Eric Alani and Michael Lichten and Valerie Borde for discussions on the manuscript.

---

**S1 Fig.**
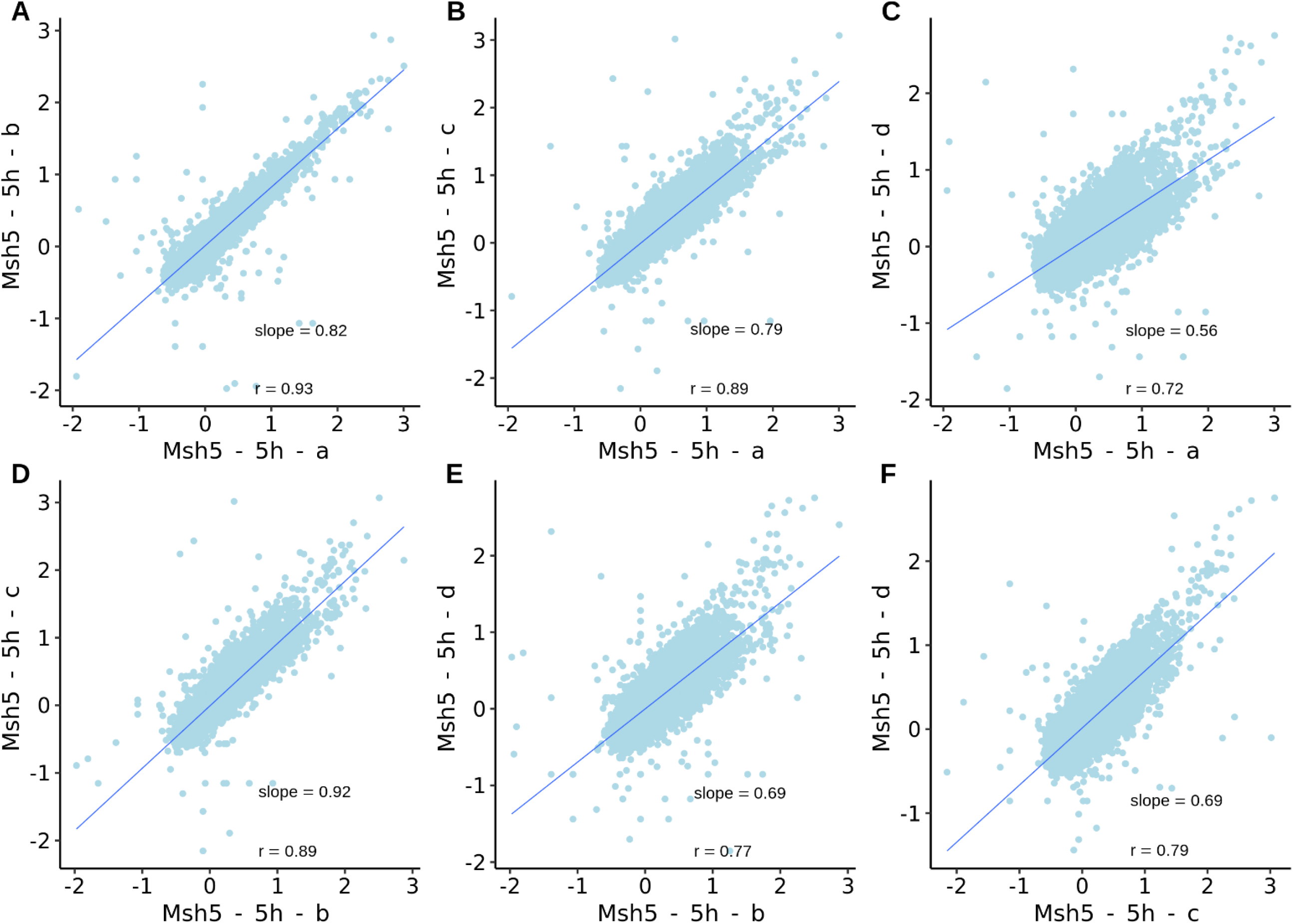
Genome-wide pairwise comparison of the log2 (fold change) of Msh5 reads for wild-type replicates (a-d) (panels A-F).

**S2 Fig.**
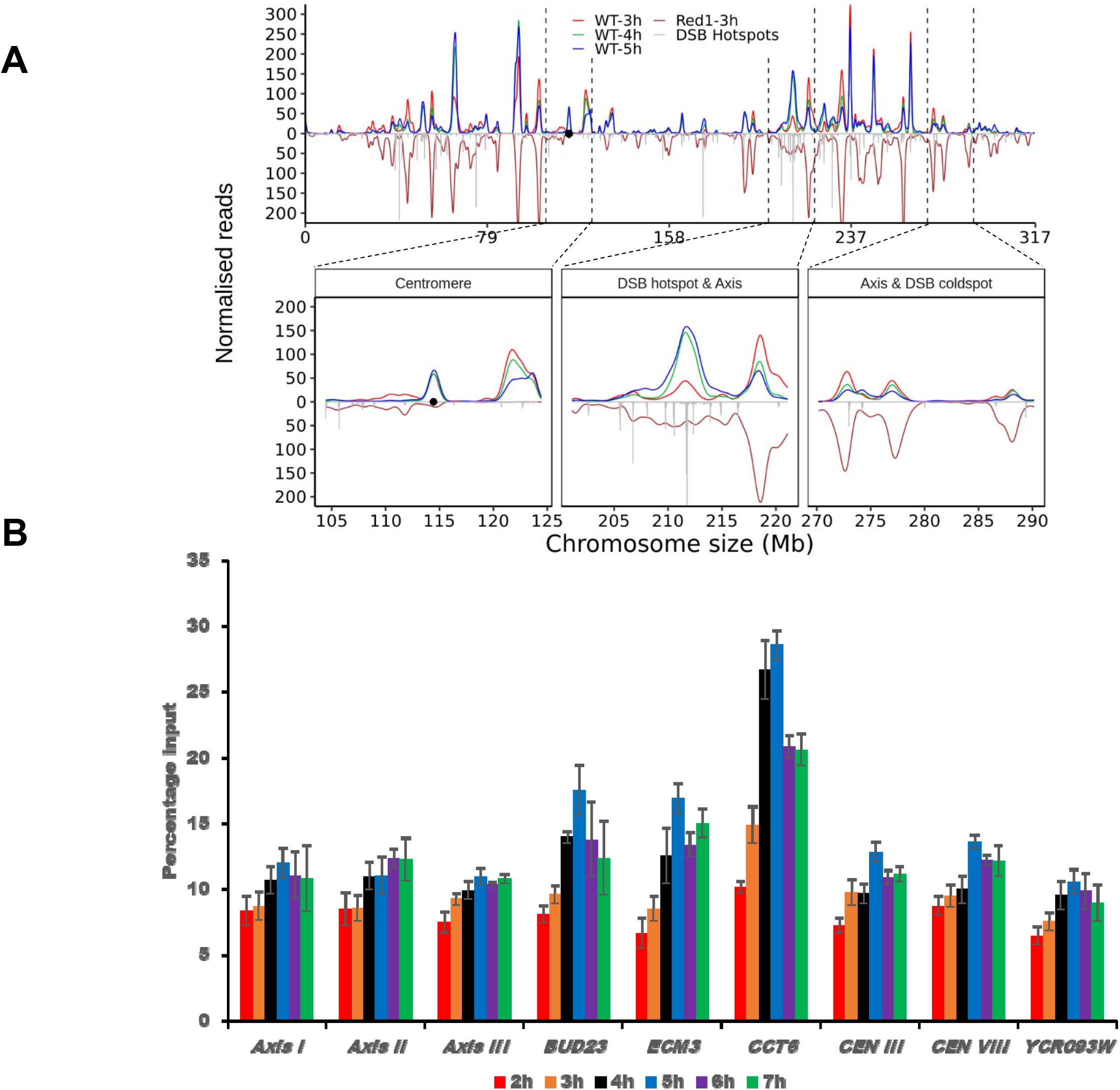
Msh5 binding sites from ChIP-Seq and ChIP-qPCR data. **A)** NCIS normalized Msh5 ChIP-Seq reads from all four replicates (not scaled with ChIP-qPCR) in wild type on chromosome III (t=3, 4, and 5h after induction of meiosis). **B)** ChIP-qPCR analysis of Msh5 enrichment at representative DSB hotspots, axis regions, centromeres, and cold spot (*YCR093W*).

**S3 Fig.**
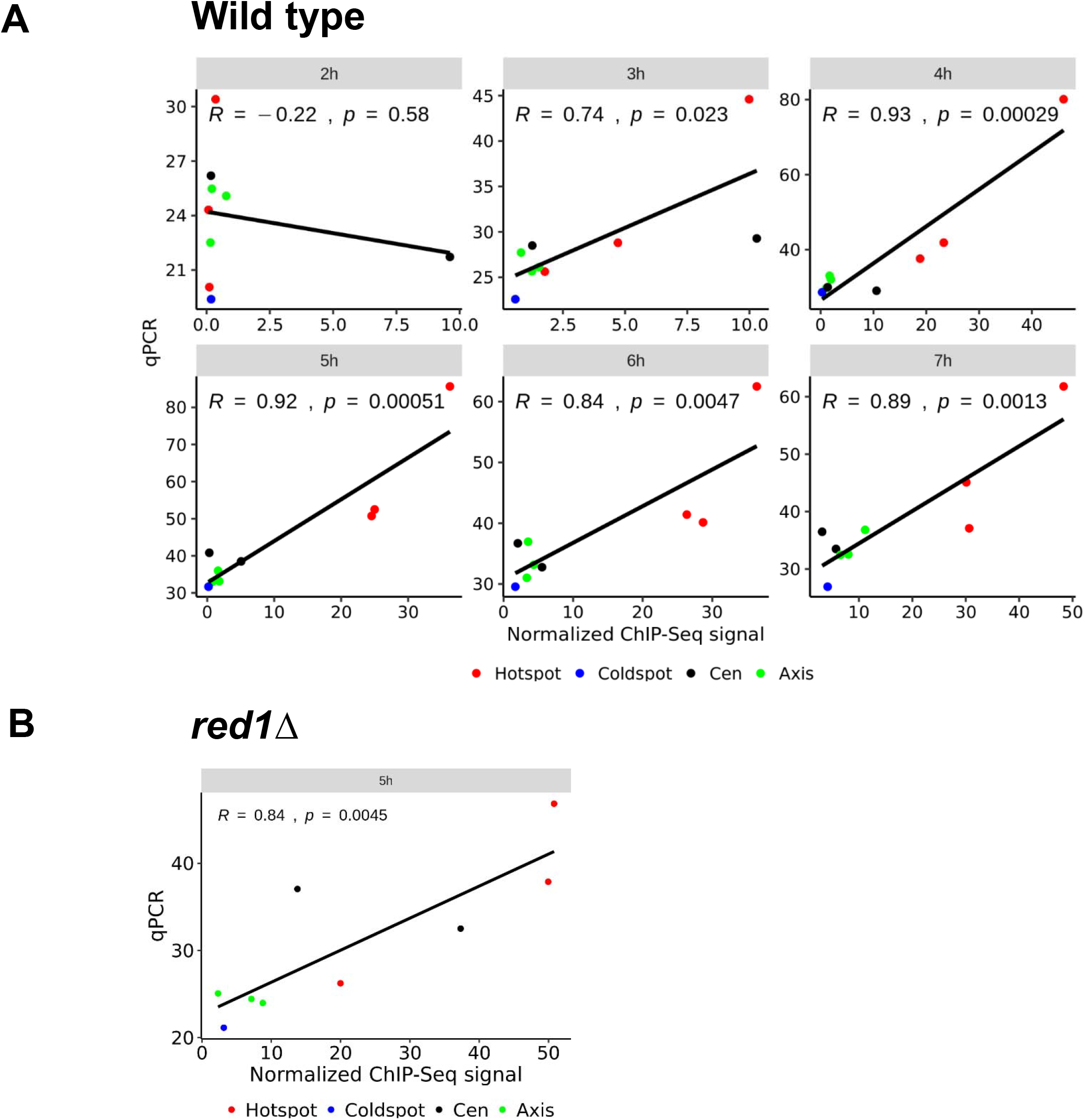
Linear regression model of Msh5 ChIP-Seq and ChIP-qPCR data. **A)** Correlation between Msh5 ChIP-qPCR and ChIP-Seq in wild-type strains at different time points (2 −7h) for DSB hotspots (*BUD23*, *ECM3*, and *CCT6*), DSB coldspot (*YCR093W*), centromere regions (*CEN I*, *CEN III*) and axis regions (*Axis I*, *Axis II*, *Axis III*). **B)** Correlation between Msh5 ChIP-qPCR and ChIP-Seq in *red1*Δ (5h).

**S4 Fig.**
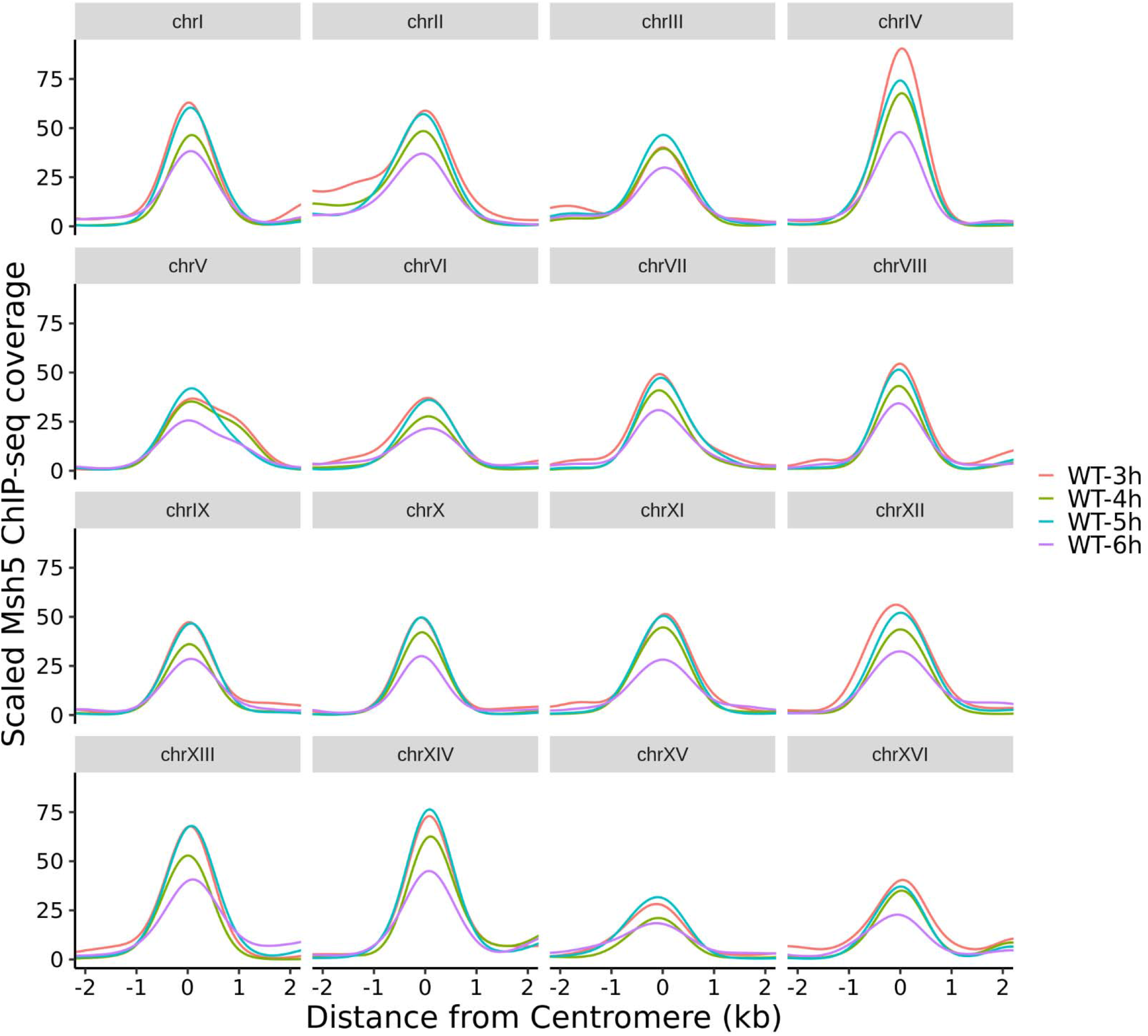
ChIP-qPCR scaled Msh5 ChIP-Seq coverage at centromere location on all chromosomes for 3-6h time points.

**S5 Fig.**
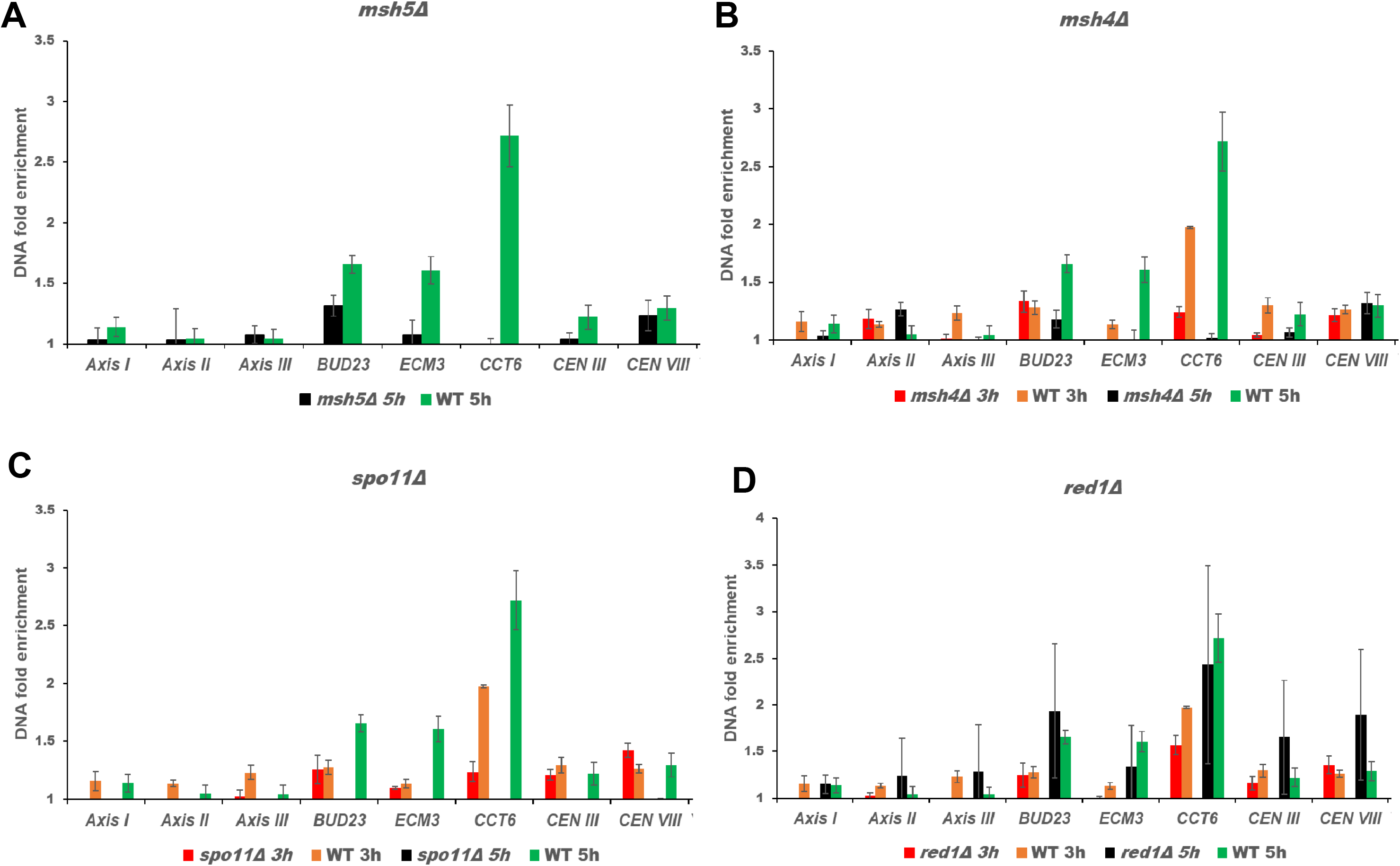
ChIP-qPCR analysis of Msh5 in meiotic mutants. ChIP-qPCR analysis of Msh5 enrichment at DSB hotspots (*BUD23*, *ECM3*, and *CCT6*), DSB coldspot (*YCR093W*), centromere regions (*CEN I, CEN III*) and axis regions (*Axis I*, *Axis II*, *Axis III*) in **A)** *msh5Δ*, **B)** *msh4Δ*, **C)** *spo11*Δ and **D)** *red1*Δ mutants. The samples are normalized using input. The error bars represent the standard deviation.

**S6 Fig.**
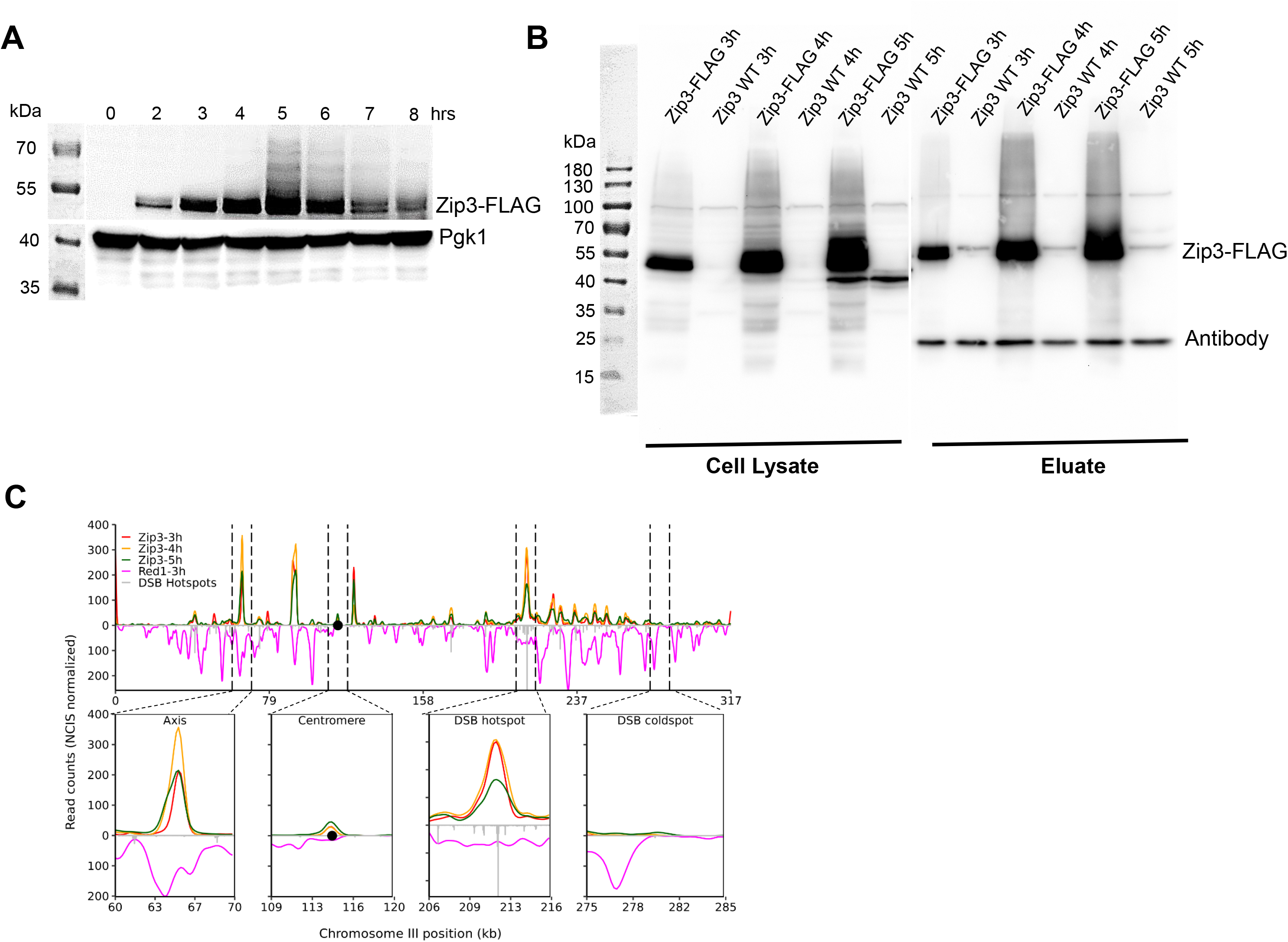
Zip3 ChIP-Seq analysis. **A)** Western blot analysis of Zip3 expression in synchronized wild-type meiosis. **B)** ChIP using anti-FLAG antibody in Zip3-FLAG and wild-type strains. Lanes 1 to 6 indicate lysate fraction at 3, 4, and 5h time points. Lanes 7 to 12 indicate eluate fractions at 3, 4, and 5h time points. **C)** Representative image of Zip3 binding on chromosome III (t=3, 4, and 5h after induction of meiosis). Zoomed in image of the centromeric region, one DSB hotspot (*YCR047C*), axis, and one DSB cold spot (*YCR093W*) is also shown.

**S7 Fig.**
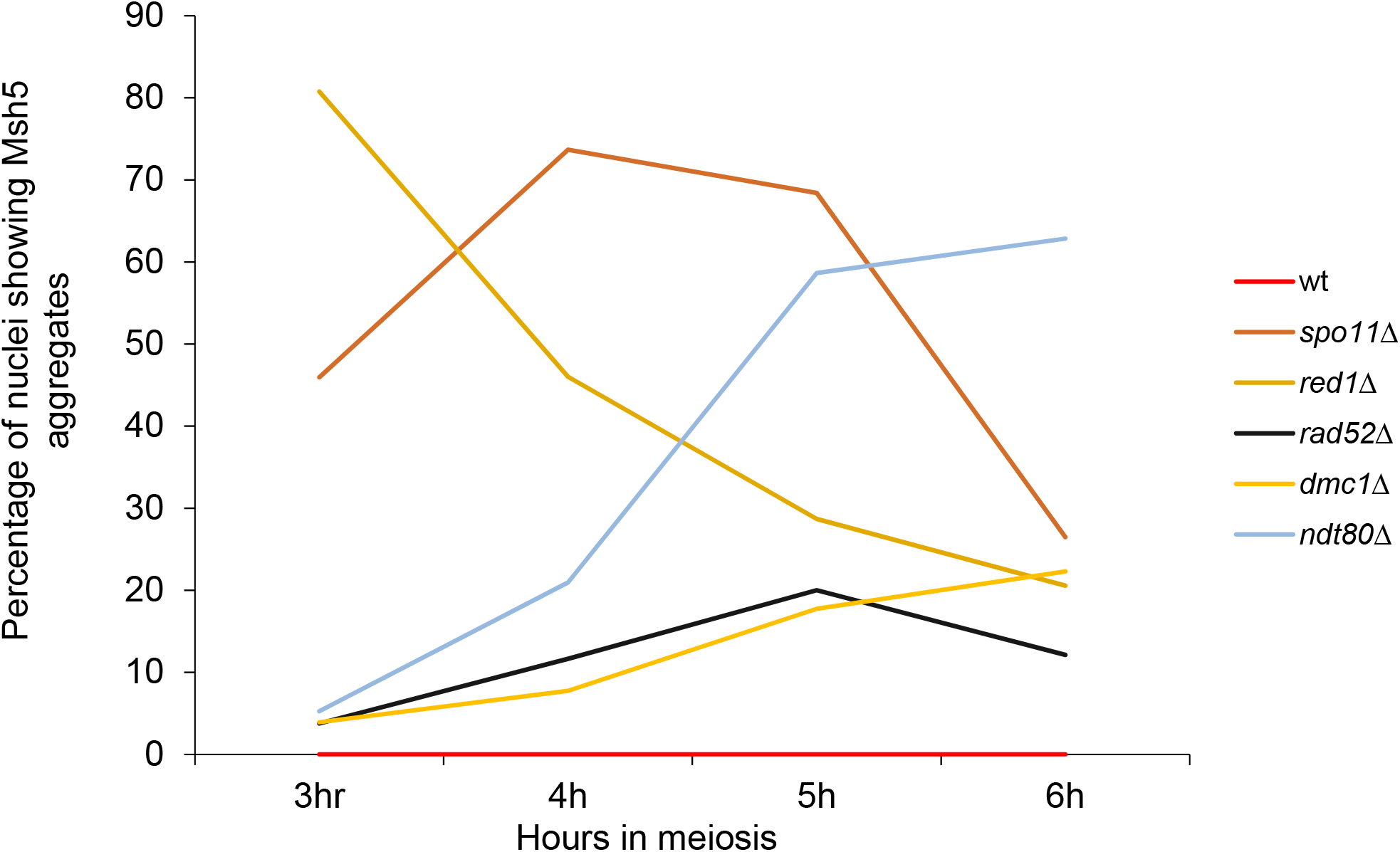
Percentage of meiotic nuclei showing Msh5 aggregate like staining. Approximately 50 nuclei, each from two independent experiments, were assessed for each time point (except *spo11Δ*, see Materials and Methods).

**S1 Table. Msh5 peaks pooled from the four wild-type replicates (WT-a,b,c,d)**. Peak start and end positions are indicated.

**S2 Table. DSB hotspots that are enriched or depleted for Msh5**. DSB hotspot data (3600 hotspots) from Pan et al. 2011[18] were sorted based on overlap with Msh5 binding regions and Spo11 oligo hits.

**S3 Table. List of strains used in this study**.

**S1 Data. Numerical data underlying all graphs in the manuscript**.

## REFERENCES

1. Neale MJ, Keeney S. Clarifying the mechanics of DNA strand exchange in meiotic recombination. Nature. 2006;442(7099):153–8. doi: 10.1038/nature04885. PubMed PMID: 16838012.

2. Keeney S, Giroux CN, Kleckner N. Meiosis-specific DNA double-strand breaks are catalyzed by Spo11, a member of a widely conserved protein family. Cell. 1997;88(3):375–84. PubMed PMID: 9039264.

3. Bergerat A, de Massy B, Gadelle D, Varoutas PC, Nicolas A, Forterre P. An atypical topoisomerase II from Archaea with implications for meiotic recombination. Nature. 1997;386(6623):414–7. doi: 10.1038/386414a0. PubMed PMID: 9121560.

4. Petronczki M, Siomos MF, Nasmyth K. Un menage a quatre: the molecular biology of chromosome segregation in meiosis. Cell. 2003;112(4):423–40. PubMed PMID: 12600308.

5. Allers T, Lichten M. Differential timing and control of noncrossover and crossover recombination during meiosis. Cell. 2001;106(1):47–57. PubMed PMID: 11461701.

6. Hunter N, Kleckner N. The single-end invasion: an asymmetric intermediate at the double-strand break to double-holliday junction transition of meiotic recombination. Cell. 2001;106(1):59–70. PubMed PMID: 11461702.

7. Page SL, Hawley RS. Chromosome choreography: the meiotic ballet. Science. 2003;301(5634):785–9. doi: 10.1126/science.1086605. PubMed PMID: 12907787.

8. Petes TD. Meiotic recombination hot spots and cold spots. Nature Reviews Genetics. 2001;2(5):360–9. doi: 10.1038/35072078. PubMed PMID: 11331902.

9. Buhler C, Borde V, Lichten M. Mapping meiotic single-strand DNA reveals a new landscape of DNA double-strand breaks in Saccharomyces cerevisiae. PLoS Biology. 2007;5(12):e324. doi: 10.1371/journal.pbio.0050324. PubMed PMID: 18076285; PubMed Central PMCID: PMC2121111.

10. Smith AV, Roeder GS. The yeast Red1 protein localizes to the cores of meiotic chromosomes. The Journal of Cell Biology. 1997;136(5):957–67. PubMed PMID: 9060462; PubMed Central PMCID: PMC2132480.

11. Zickler D, Kleckner N. A few of our favorite things: Pairing, the bouquet, crossover interference and evolution of meiosis. Seminars in Cell & Developmental Biology. 2016;54:135–48. doi: 10.1016/j.semcdb.2016.02.024. PubMed PMID: 26927691; PubMed Central PMCID: PMC4867269.

12. Zickler D, Kleckner N. Meiotic chromosomes: integrating structure and function. Annual Review of Genetics. 1999;33:603–754. doi: 10.1146/annurev.genet.33.1.603. PubMed PMID: 10690419.

13. Hollingsworth NM, Goetsch L, Byers B. The HOP1 gene encodes a meiosis-specific component of yeast chromosomes. Cell. 1990;61(1):73–84. PubMed PMID: 2107981.

14. Klein F, Mahr P, Galova M, Buonomo SB, Michaelis C, Nairz K, et al. A central role for cohesins in sister chromatid cohesion, formation of axial elements, and recombination during yeast meiosis. Cell. 1999;98(1):91–103. doi: 10.1016/S0092-8674(00)80609-1. PubMed PMID: 10412984.

15. Sun X, Huang L, Markowitz TE, Blitzblau HG, Chen D, Klein F, et al. Transcription dynamically patterns the meiotic chromosome-axis interface. eLife. 2015;4. doi: 10.7554/eLife.07424. PubMed PMID: 26258962; PubMed Central PMCID: PMC4530585.

16. Blat Y, Protacio RU, Hunter N, Kleckner N. Physical and functional interactions among basic chromosome organizational features govern early steps of meiotic chiasma formation. Cell. 2002;111(6):791–802. PubMed PMID: 12526806.

17. Borde V, de Massy B. Programmed induction of DNA double strand breaks during meiosis: setting up communication between DNA and the chromosome structure. Current opinion in Genetics & Development. 2013;23(2):147–55. doi: 10.1016/j.gde.2012.12.002. PubMed PMID: 23313097.

18. Pan J, Sasaki M, Kniewel R, Murakami H, Blitzblau HG, Tischfield SE, et al. A hierarchical combination of factors shapes the genome-wide topography of yeast meiotic recombination initiation. Cell. 2011;144(5):719–31. doi: 10.1016/j.cell.2011.02.009. PubMed PMID: 21376234; PubMed Central PMCID: PMCPMC3063416.

19. Panizza S, Mendoza MA, Berlinger M, Huang L, Nicolas A, Shirahige K, et al. Spo11-accessory proteins link double-strand break sites to the chromosome axis in early meiotic recombination. Cell. 2011;146(3):372–83. doi: 10.1016/j.cell.2011.07.003. PubMed PMID: 21816273.

20. Kleckner N. Chiasma formation: chromatin/axis interplay and the role(s) of the synaptonemal complex. Chromosoma. 2006;115(3):175–94. doi: 10.1007/s00412-006-0055-7. PubMed PMID: 16555016.

21. Acquaviva L, Szekvolgyi L, Dichtl B, Dichtl BS, de La Roche Saint Andre C, Nicolas A, et al. The COMPASS subunit Spp1 links histone methylation to initiation of meiotic recombination. Science. 2013;339(6116):215–8. doi: 10.1126/science.1225739. PubMed PMID: 23160953.

22. Sommermeyer V, Beneut C, Chaplais E, Serrentino ME, Borde V. Spp1, a member of the Set1 Complex, promotes meiotic DSB formation in promoters by tethering histone H3K4 methylation sites to chromosome axes. Molecular Cell. 2013;49(1):43–54. doi: 10.1016/j.molcel.2012.11.008. PubMed PMID: 23246437.

23. Medhi D, Goldman AS, Lichten M. Local chromosome context is a major determinant of crossover pathway biochemistry during budding yeast meiosis. eLife. 2016;5. doi: 10.7554/eLife.19669. PubMed PMID: 27855779; PubMed Central PMCID: PMC5222560.

24. Borner GV, Kleckner N, Hunter N. Crossover/noncrossover differentiation, synaptonemal complex formation, and regulatory surveillance at the leptotene/zygotene transition of meiosis. Cell. 2004;117(1):29–45. PubMed PMID: 15066280.

25. Shinohara M, Oh SD, Hunter N, Shinohara A. Crossover assurance and crossover interference are distinctly regulated by the ZMM proteins during yeast meiosis. Nature Genetics. 2008;40(3):299–309. doi: 10.1038/ng.83. PubMed PMID: 18297071.

26. Lynn A, Soucek R, Borner GV. ZMM proteins during meiosis: crossover artists at work. Chromosome Research. 2007;15(5):591–605. doi: 10.1007/s10577-007-1150-1. PubMed PMID: 17674148.

27. Manhart CM, Ni X, White MA, Ortega J, Surtees JA, Alani E. The mismatch repair and meiotic recombination endonuclease Mlh1-Mlh3 is activated by polymer formation and can cleave DNA substrates in trans. PLoS Biology. 2017;15(4):e2001164. doi: 10.1371/journal.pbio.2001164. PubMed PMID: 28453523; PubMed Central PMCID: PMC5409509.

28. Zakharyevich K, Ma Y, Tang S, Hwang PY, Boiteux S, Hunter N. Temporally and biochemically distinct activities of Exo1 during meiosis: double-strand break resection and resolution of double Holliday junctions. Molecular Cell. 2010;40(6):1001–15. doi: 10.1016/j.molcel.2010.11.032. PubMed PMID: 21172664; PubMed Central PMCID: PMC3061447.

29. de los Santos T, Hunter N, Lee C, Larkin B, Loidl J, Hollingsworth NM. The Mus81/Mms4 endonuclease acts independently of double-Holliday junction resolution to promote a distinct subset of crossovers during meiosis in budding yeast. Genetics. 2003;164(1):81–94. PubMed PMID: 12750322; PubMed Central PMCID: PMCPMC1462551.

30. Argueso JL, Wanat J, Gemici Z, Alani E. Competing crossover pathways act during meiosis in Saccharomyces cerevisiae. Genetics. 2004;168(4):1805–16. doi: 10.1534/genetics.104.032912. PubMed PMID: 15611158; PubMed Central PMCID: PMCPMC1448724.

31. Ross-Macdonald P, Roeder GS. Mutation of a meiosis-specific MutS homolog decreases crossing over but not mismatch correction. Cell. 1994;79(6):1069–80. PubMed PMID: 8001134.

32. Pochart P, Woltering D, Hollingsworth NM. Conserved properties between functionally distinct MutS homologs in yeast. J Biol Chem. 1997;272(48):30345–9. doi: DOI 10.1074/jbc.272.48.30345. PubMed PMID: WOS:A1997YH61300055.

33. Hollingsworth NM, Ponte L, Halsey C. MSH5, a novel MutS homolog, facilitates meiotic reciprocal recombination between homologs in Saccharomyces cerevisiae but not mismatch repair. Genes & Development. 1995;9(14):1728–39. PubMed PMID: 7622037.

34. Snowden T, Acharya S, Butz C, Berardini M, Fishel R. hMSH4-hMSH5 recognizes Holliday Junctions and forms a meiosis-specific sliding clamp that embraces homologous chromosomes. Molecular Cell. 2004;15(3):437–51. doi: 10.1016/j.molcel.2004.06.040. PubMed PMID: 15304223.

35. Lahiri S, Li Y, Hingorani MM, Mukerji I. MutSgamma-Induced DNA conformational changes provide insights into its role in meiotic recombination. Biophysical Journal. 2018;115(11):2087–101. Epub 2018/11/24. doi: 10.1016/j.bpj.2018.10.029. PubMed PMID: 30467025; PubMed Central PMCID: PMCPMC6289823.

36. He W, Rao H, Tang S, Bhagwat N, Kulkarni DS, Ma Y, et al. Regulated proteolysis of MutSgamma controls meiotic crossing over. Molecular Cell. 2020;78(1):168–83 e5. Epub 2020/03/05. doi: 10.1016/j.molcel.2020.02.001. PubMed PMID: 32130890.

37. Nishant KT, Plys AJ, Alani E. A mutation in the putative MLH3 endonuclease domain confers a defect in both mismatch repair and meiosis in Saccharomyces cerevisiae. Genetics. 2008;179(2):747–55. doi: 10.1534/genetics.108.086645. PubMed PMID: 18505871; PubMed Central PMCID: PMCPMC2429871.

38. Zakharyevich K, Tang S, Ma Y, Hunter N. Delineation of joint molecule resolution pathways in meiosis identifies a crossover-specific resolvase. Cell. 2012;149(2):334–47. doi: 10.1016/j.cell.2012.03.023. PubMed PMID: 22500800; PubMed Central PMCID: PMCPMC3377385.

39. Kolas NK, Svetlanov A, Lenzi ML, Macaluso FP, Lipkin SM, Liskay RM, et al. Localization of MMR proteins on meiotic chromosomes in mice indicates distinct functions during prophase I. The Journal of Cell Biology. 2005;171(3):447–58. doi: 10.1083/jcb.200506170. PubMed PMID: 16260499; PubMed Central PMCID: PMC2171243.

40. Cannavo E, Sanchez A, Anand R, Ranjha L, Hugener J, Adam C, et al. Regulation of the MLH1-MLH3 endonuclease in meiosis. 2020:2020.02.12.946293. doi: 10.1101/2020.02.12.946293%JbioRxiv.

41. Kulkarni DS, Owens S, Honda M, Ito M, Yang Y, Corrigan MW, et al. PCNA activates the MutLγ endonuclease to promote meiotic crossing over. 2020:2020.02.12.946020. doi: 10.1101/2020.02.12.946020%JbioRxiv.

42. Kneitz B, Cohen PE, Avdievich E, Zhu L, Kane MF, Hou H, Jr., et al. MutS homolog 4 localization to meiotic chromosomes is required for chromosome pairing during meiosis in male and female mice. Genes & Development. 2000;14(9):1085–97. PubMed PMID: 10809667; PubMed Central PMCID: PMC316572.

43. Edelmann W, Cohen PE, Kneitz B, Winand N, Lia M, Heyer J, et al. Mammalian MutS homologue 5 is required for chromosome pairing in meiosis. Nature Genetics. 1999;21(1):123–7. doi: 10.1038/5075. PubMed PMID: 9916805.

44. Cole F, Kauppi L, Lange J, Roig I, Wang R, Keeney S, et al. Homeostatic control of recombination is implemented progressively in mouse meiosis. Nature Cell Biology. 2012;14(4):424–30. doi: 10.1038/ncb2451. PubMed PMID: 22388890; PubMed Central PMCID: PMC3319518.

45. Svetlanov A, Cohen PE. Mismatch repair proteins, meiosis, and mice: understanding the complexities of mammalian meiosis. Experimental Cell Research. 2004;296(1):71–9. doi: 10.1016/j.yexcr.2004.03.020. PubMed PMID: 15120996.

46. Santucci-Darmanin S, Walpita D, Lespinasse F, Desnuelle C, Ashley T, Paquis-Flucklinger V. MSH4 acts in conjunction with MLH1 during mammalian meiosis. FASEB Journal. 2000;14(11):1539–47. PubMed PMID: 10928988.

47. De Muyt A, Pyatnitskaya A, Andreani J, Ranjha L, Ramus C, Laureau R, et al. A meiotic XPF-ERCC1-like complex recognizes joint molecule recombination intermediates to promote crossover formation. Genes & Development. 2018;32(3-4):283–96. doi: 10.1101/gad.308510.117. PubMed PMID: 29440262; PubMed Central PMCID: PMCPMC5859969.

48. Serrentino ME, Chaplais E, Sommermeyer V, Borde V. Differential association of the conserved SUMO ligase Zip3 with meiotic double-strand break sites reveals regional variations in the outcome of meiotic recombination. PLoS Genetics. 2013;9(4):e1003416. doi: 10.1371/journal.pgen.1003416. PubMed PMID: 23593021; PubMed Central PMCID: PMCPMC3616913.

49. Shinohara M, Hayashihara K, Grubb JT, Bishop DK, Shinohara A. DNA damage response clamp 9-1-1 promotes assembly of ZMM proteins for formation of crossovers and synaptonemal complex. Journal of Cell Science. 2015;128(8):1494–506. doi: 10.1242/jcs.161554. PubMed PMID: 25736290; PubMed Central PMCID: PMCPMC4518444.

50. Nishant KT, Chen C, Shinohara M, Shinohara A, Alani E. Genetic analysis of baker’s yeast Msh4-Msh5 reveals a threshold crossover level for meiotic viability. PLoS Genetics. 2010;6(8). doi: 10.1371/journal.pgen.1001083. PubMed PMID: 20865162; PubMed Central PMCID: PMCPMC2928781.

51. Mimitou EP, Yamada S, Keeney S. A global view of meiotic double-strand break end resection. Science. 2017;355(6320):40–5. doi: 10.1126/science.aak9704. PubMed PMID: 28059759; PubMed Central PMCID: PMC5234563.

52. Bishop DK, Park D, Xu L, Kleckner N. DMC1: a meiosis-specific yeast homolog of E. coli recA required for recombination, synaptonemal complex formation, and cell cycle progression. Cell. 1992;69(3):439–56. PubMed PMID: 1581960.

53. Lao JP, Cloud V, Huang CC, Grubb J, Thacker D, Lee CY, et al. Meiotic crossover control by concerted action of Rad51-Dmc1 in homolog template bias and robust homeostatic regulation. PLoS Genetics. 2013;9(12):e1003978. doi: 10.1371/journal.pgen.1003978. PubMed PMID: 24367271; PubMed Central PMCID: PMCPMC3868528.

54. Cloud V, Chan YL, Grubb J, Budke B, Bishop DK. Rad51 is an accessory factor for Dmc1-mediated joint molecule formation during meiosis. Science. 2012;337(6099):1222–5. doi: 10.1126/science.1219379. PubMed PMID: 22955832; PubMed Central PMCID: PMCPMC4056682.

55. Lao JP, Oh SD, Shinohara M, Shinohara A, Hunter N. Rad52 promotes postinvasion steps of meiotic double-strand-break repair. Molecular Cell. 2008;29(4):517–24. doi: 10.1016/j.molcel.2007.12.014. PubMed PMID: 18313389; PubMed Central PMCID: PMC2351957.

56. Bishop DK. RecA homologs Dmc1 and Rad51 interact to form multiple nuclear complexes prior to meiotic chromosome synapsis. Cell. 1994;79(6):1081–92. PubMed PMID: 7528104.

57. Murakami H, Lam I, Huang PC, Song J, van Overbeek M, Keeney S. Multilayered mechanisms ensure that short chromosomes recombine in meiosis. Nature. 2020;582(7810):124–8. Epub 2020/06/05. doi: 10.1038/s41586-020-2248-2. PubMed PMID: 32494071; PubMed Central PMCID: PMCPMC7298877.

58. Chakraborty P, Pankajam AV, Lin G, Dutta A, Krishnaprasad GN, Tekkedil MM, et al. Modulating crossover frequency and interference for obligate crossovers in Saccharomyces cerevisiae meiosis. G3 (Bethesda). 2017;7(5):1511–24. doi: 10.1534/g3.117.040071. PubMed PMID: 28315832; PubMed Central PMCID: PMCPMC5427503.

59. Krishnaprasad GN, Anand MT, Lin G, Tekkedil MM, Steinmetz LM, Nishant KT. Variation in crossover frequencies perturb crossover assurance without affecting meiotic chromosome segregation in Saccharomyces cerevisiae. Genetics. 2015;199(2):399–412. Epub 2014/12/04. doi: 10.1534/genetics.114.172320. PubMed PMID: 25467183; PubMed Central PMCID: PMCPMC4317650.

60. Prieler S, Penkner A, Borde V, Klein F. The control of Spo11’s interaction with meiotic recombination hotspots. Genes & Development. 2005;19(2):255–69. doi: 10.1101/gad.321105. PubMed PMID: 15655113; PubMed Central PMCID: PMC545890.

61. Schwacha A, Kleckner N. Interhomolog bias during meiotic recombination: meiotic functions promote a highly differentiated interhomolog-only pathway. Cell. 1997;90(6):1123–35. PubMed PMID: 9323140.

62. Carballo JA, Johnson AL, Sedgwick SG, Cha RS. Phosphorylation of the axial element protein Hop1 by Mec1/Tel1 ensures meiotic interhomolog recombination. Cell. 2008;132(5):758–70. doi: 10.1016/j.cell.2008.01.035. PubMed PMID: 18329363.

63. Kim KP, Weiner BM, Zhang L, Jordan A, Dekker J, Kleckner N. Sister cohesion and structural axis components mediate homolog bias of meiotic recombination. Cell. 2010;143(6):924–37. doi: 10.1016/j.cell.2010.11.015. PubMed PMID: 21145459; PubMed Central PMCID: PMC3033573.

64. Evans E, Sugawara N, Haber JE, Alani E. The Saccharomyces cerevisiae Msh2 mismatch repair protein localizes to recombination intermediates in vivo. Molecular Cell. 2000;5(5):789–99. PubMed PMID: 10882115.

65. Agarwal S, Roeder GS. Zip3 provides a link between recombination enzymes and synaptonemal complex proteins. Cell. 2000;102(2):245–55. PubMed PMID: 10943844.

66. Pyatnitskaya A, Borde V, De Muyt A. Crossing and zipping: molecular duties of the ZMM proteins in meiosis. Chromosoma. 2019;128(3):181–98. Epub 2019/06/27. doi: 10.1007/s00412-019-00714-8. PubMed PMID: 31236671.

67. Tsubouchi T, Macqueen AJ, Roeder GS. Initiation of meiotic chromosome synapsis at centromeres in budding yeast. Genes & Development. 2008;22(22):3217–26. Epub 2008/12/06. doi: 10.1101/gad.1709408. PubMed PMID: 19056898; PubMed Central PMCID: PMCPMC2593611.

68. Hunter N, Borts RH. Mlh1 is unique among mismatch repair proteins in its ability to promote crossing-over during meiosis. Genes & Development. 1997;11(12):1573–82. Epub 1997/06/15. doi: 10.1101/gad.11.12.1573. PubMed PMID: 9203583.

69. Kleckner N, Zickler D, Jones GH, Dekker J, Padmore R, Henle J, et al. A mechanical basis for chromosome function. Proceedings of the National Academy of Sciences of the United States of America. 2004;101(34):12592–7. doi: 10.1073/pnas.0402724101. PubMed PMID: 15299144; PubMed Central PMCID: PMC515102.

70. Zhang L, Wang S, Yin S, Hong S, Kim KP, Kleckner N. Topoisomerase II mediates meiotic crossover interference. Nature. 2014;511(7511):551–6. doi: 10.1038/nature13442. PubMed PMID: 25043020; PubMed Central PMCID: PMCPMC4128387.

71. Mortimer RK, Johnston JR. Genealogy of principal strains of the yeast genetic stock center. Genetics. 1986;113(1):35–43. PubMed PMID: 3519363; PubMed Central PMCID: PMC1202798.

72. Rose MD, F. Winston, and P. Hieter. Methods in yeast genetics. Cold Spring Harbor Laboratory Press, Cold Spring Harbor, NY. 1990.

73. McCusker JH, Clemons KV, Stevens DA, Davis RW. Genetic characterization of pathogenic Saccharomyces cerevisiae isolates. Genetics. 1994;136(4):1261–9. PubMed PMID: 8013903; PubMed Central PMCID: PMC1205906.

74. Gietz RD, Schiestl RH, Willems AR, Woods RA. Studies on the transformation of intact yeast cells by the LiAc/SS-DNA/PEG procedure. Yeast. 1995;11(4):355–60. doi: 10.1002/yea.320110408. PubMed PMID: 7785336.

75. Goldstein AL, McCusker JH. Three new dominant drug resistance cassettes for gene disruption in Saccharomyces cerevisiae. Yeast. 1999;15(14):1541–53. doi: 10.1002/(SICI)1097-0061(199910)15:14<1541::AID-YEA476>3.0.CO;2-K. PubMed PMID: 10514571.

76. Thacker D, Mohibullah N, Zhu X, Keeney S. Homologue engagement controls meiotic DNA break number and distribution. Nature. 2014;510(7504):241–6. doi: 10.1038/nature13120. PubMed PMID: 24717437; PubMed Central PMCID: PMC4057310.

77. Mendoza MA, Panizza S, Klein F. Analysis of protein-DNA interactions during meiosis by quantitative chromatin immunoprecipitation (qChIP). Methods in Molecular Biology. 2009;557:267–83. doi: 10.1007/978-1-59745-527-5_17. PubMed PMID: 19799188.

78. Liang K, Keles S. Normalization of ChIP-seq data with control. BMC Bioinformatics. 2012;13:199. doi: 10.1186/1471-2105-13-199. PubMed PMID: 22883957; PubMed Central PMCID: PMC3475056.

79. Zhang Y, Liu T, Meyer CA, Eeckhoute J, Johnson DS, Bernstein BE, et al. Model-based analysis of ChIP-Seq (MACS). Genome Biology. 2008;9(9):R137. doi: 10.1186/gb-2008-9-9-r137. PubMed PMID: 18798982; PubMed Central PMCID: PMC2592715.

80. Murakami H, Keeney S. Temporospatial coordination of meiotic DNA replication and recombination via DDK recruitment to replisomes. Cell. 2014;158(4):861–73. Epub 2014/08/16. doi: 10.1016/j.cell.2014.06.028. PubMed PMID: 25126790; PubMed Central PMCID: PMCPMC4141489.

