## Supplementary material for "The bakers’s yeast Msh4-Msh5 associates with double-strand break hotspots and chromosome axis during meiosis to promote crossovers": S3 Table

**S3 Table. List of strains used in this study**

| **Strain** | **Genotype** | **Source** |
| --- | --- | --- |
| NHY1162 | *MATα ho::hisG,leu2::hisG,ura3(∆sma-pst),his4-X::LEU2-(NgoM IV-URA3)* | Martini, E *et al*. 2006 |
| KTY 629 | as NHY1162 except *spo11∆::kanMX4* | This study |
| KTY 633 | as NHY1162 except *dmc1∆::kanMX4* | This study |
| KTY 637 | as NHY1162 except *rad52∆::kanMX4* | This study |
| KTY 641 | as NHY1162 except *ndt80∆::kanMX4* | This study |
| KTY 645 | as NHY1162 except *red1∆::kanMX4* | This study |
| KTY 715 | as NHY1162 except *msh4∆::natMX4* | This study |
| KTY 719 | as NHY1162 except *msh5∆::natMX4* | This study |
| KTY 733 | as NHY1162 except *ZIP3-His6-Flag3::kanMX4* | This study |
| NHY1168 | *MATa, ho::hisG, leu::hisG, ura3(∆Sma-Pst), HIS4::LEU2-(BamHI)* | Martini, E *et al*. 2006 |
| KTY 631 | as NHY1168 except *spo11∆::kanMX4* | This study |
| KTY 635 | as NHY1168 except *dmc1∆::kanMX4* | This study |
| KTY 639 | as NHY1168 except *rad52∆::kanMX4* | This study |
| KTY 643 | as NHY1168 except *ndt80∆::kanMX4* | This study |
| KTY 647 | as NHY1168 except *red1∆::kanMX4* | This study |
| KTY 713 | as NHY1168 except *msh4∆::natMX4* | This study |
| KTY 717 | as NHY1168 except *msh5∆::natMX4* | This study |
| KTY 734 | as NHY1168 except *ZIP3-His6-Flag3::kanMX4* | This study |

Martini E, Diaz RL, Hunter N, Keeney, S. Crossover homeostasis in yeast meiosis. Cell 2006; 126: 285-295.
